# Tuning inhibitor susceptibility of human 4-hydroxyphenylpyruvate dioxygenase to rescue tyrosine catabolism in the presence of β-triketones

**DOI:** 10.64898/2026.10.22.760921

**Authors:** Jessie Neuckermans, Alan Mertens, Sien Lequeue, Nina S. Salvi, Liesbeth Desmet, Gigly G. Del’Haye, Karina Tretowicz, Tamara Vanhaecke, Ulrich Schwaneberg, Joery De Kock

## Abstract

Human 4-hydroxyphenylpyruvate dioxygenase (HPD) catalyses a key step in tyrosine degradation via homogentisate in liver and kidney cells. Pharmacological inhibition of HPD with nitisinone (NTBC) is standard treatment for tyrosinemia type 1 (HT1) and alkaptonuria (AKU). Although life-changing, continuous NTBC treatment limits metabolic flexibility and complicates emerging gene- and cell-based therapeutic approaches. Engineering HPD variants that retain catalytic activity during NTBC exposure could facilitate restoration of tyrosine metabolism, yet the molecular determinants of inhibitor tolerance remain poorly understood. Here, we applied Knowledge Gaining Directed Evolution (KnowVolution) to enhance NTBC tolerance while considering enzyme activity, predicted stability, and MHC-II peptide binding. Random mutagenesis, site-saturation mutagenesis, and computational analyses identified beneficial substitutions that improved enzyme activity in the presence of NTBC. These substitutions are predominantly located outside the catalytic cavity, suggesting that NTBC tolerance may involve distal structural effects rather than direct modification of inhibitor-binding residues. Recombination of selected substitutions, guided by activity and computational predictions of stability and MHC-II binding, yielded multi-substitution HPD variants with markedly increased NTBC tolerance. The leading variant, HPD-M8, exhibited up to 40-fold greater NTBC tolerance than wild-type HPD and increased tolerance to structurally related β-triketone inhibitors, without an overall increase in predicted MHC-II binder burden. Computational analyses suggested altered catalytic access-pathway energetics without major perturbation of the conserved catalytic machinery, consistent with a possible contribution of distal substitutions to inhibitor tolerance. Together, these findings provide insight into HPD inhibitor tolerance and establish a foundation for future HT1 and AKU therapeutic strategies.

## 1. Introduction

Human 4-hydroxyphenylpyruvate dioxygenase (HPD; EC 1.13.11.27) is a non-heme iron (II)-dependent oxygenase that catalyses the second step in tyrosine degradation: the conversion of 4-hydroxyphenylpyruvate (HPP) to homogentisate (HGA) (Fig. 1a), with concomitant CO_2_ release. This complex reaction, involving oxidative decarboxylation, aromatic hydroxylation, and side-chain migration within a single catalytic cycle, is essential for funnelling tyrosine toward fumarate and acetoacetate[1–4].

**Fig. 1:**
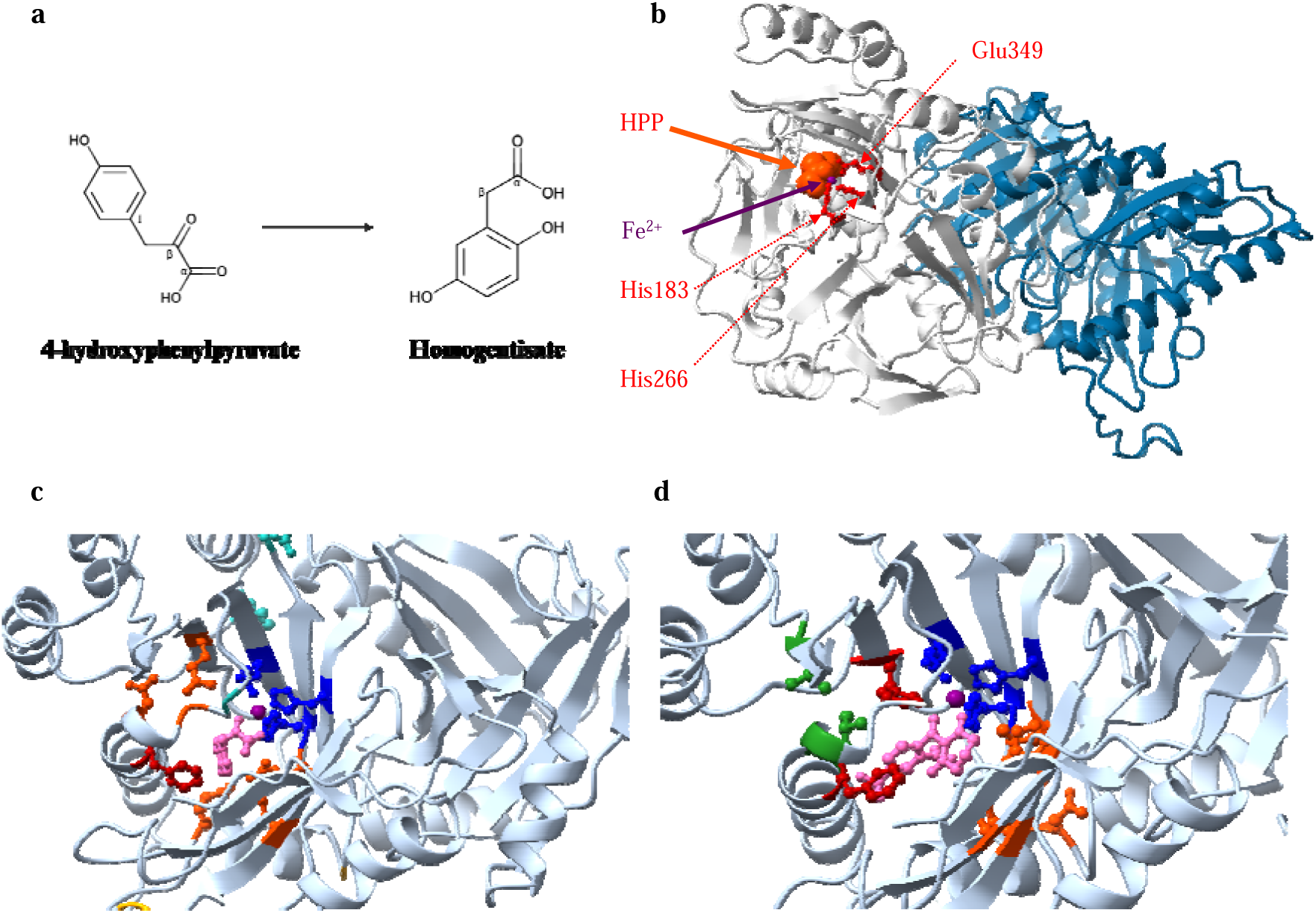
Structure and catalytic mechanism of human HPD. (a) Chemical conversion of HPP to HGA, catalysed by HPD (structures obtained from ChemSpider **-**Created in BioRender. Lequeue, S. (2026) https://BioRender.com/ujgkc2c). (b) Homodimeric structure of human HPD shown as cartoon representation with chain A in grey and chain B in blue. The catalytic Fe² ion (purple sphere) is coordinated by the conserved 2-His-1-carboxylate motif consisting of His183, His266, and Glu349 (red). (c) Close-up view of the catalytic cavity showing key residues involved in substrate coordination and catalysis. HPP is shown in pink and the catalytic Fe² ion in purple. Residues (His183, His266, Glu349) coordinating the Fe² ion are shown in dark blue as ball-and-stick representations. The carboxyl group of HPP forms a hydrogen bond with Gln334, Asn363 (shown in orange) and Glu349 (shown in cyan blue), the 4-hydroxyl group on the aromatic ring interacts with Gln251 and Gln265 (shown in orange), π-π interactions essential for the positioning of the benzene ring of the HPP for oxidation involve Phe364 (shown in red). Gln272, Gln286, and Gln358 (shown in green) play a central role in HPP binding and the first nucleophilic attack. Asn261 and Ser246 (shown in yellow) play a key role in the electrophilic attack of the aromatic C1 for hydroxylation and the final ortho-rearrangement [8]. (d) Binding mode of the inhibitor NTBC (pink) within the catalytic cavity. Two carbonyl groups of NTBC are conjugated with the Fe^2+^, which is also chelated by the conservative triad of His183, His266, and Glu349 (shown in dark blue as ball and sticks). The cyclohexanedione is surrounded by Val185, Pro239, Asn241, Ser226, and Phe 259 (shown in orange), the phenyl moiety of NTBC forms π-π interactions with Phe336 and Phe364 (shown in red), and the trifluoromethyl group is bound to Phe336, Phe364 (shown in red), and Leu323, Asn362, and Leu367 (shown in green) [7]. Structural figures were generated using UCSF ChimeraX.

Structurally, human HPD is a 393-residue homodimer with a conserved C-terminal β-barrel that houses the catalytic Fe(II), coordinated by the canonical 2-His-1-carboxylate motif (His183, His266, Glu349) (Fig. 1b-c). Access to this buried catalytic core is regulated by a C-terminal α-helix that functions as a gating element [2,5,6]. Recent structural and computational studies further demonstrated that ligand binding induces pronounced conformational rearrangements of the C-terminal gating helix and adjacent loop regions, stabilizing an open state of the catalytic cavity and facilitating substrate or inhibitor accommodation [7–9]. Despite limited sequence identity across species, the structural fold is highly conserved, and mutagenesis studies have highlighted the contribution of residues throughout both N- and C-termini for catalysis and stability [2,3,5,6,10].

Loss of HPD function leads to upstream metabolite accumulation and causes the metabolic disorder hereditary tyrosinemia type 3, characterized by neurocognitive impairment, while specific dominant HPD mutations, e.g. c.722A>G (p.N241S) and c.97G>A (p.A33T), exist that cause hawkinsinuria, an ultra-rare disorder characterized by renal excretion of the unusual tyrosine metabolites hawkinsin and 4-hydroxycyclohexylacetic acid [1,11–14]. However, HPD is the main therapeutic target in hereditary tyrosinemia type 1 (HT1) and alkaptonuria (AKU). In both diseases, pharmacological HPD inhibition reduces HGA formation and prevents accumulation of downstream toxic intermediates [4,15,16]. The only approved HPD inhibitor, nitisinone or 2-(2-nitro-4-trifluoromethylbenzoyl)-1,3-cyclohexanedione (NTBC), is a potent β-triketone originally developed as a herbicide [17,18]. Structurally mimicking HPP, NTBC forms a tight, slowly dissociating enzyme–inhibitor complex through reversible π-stacking interactions with active-site phenylalanine residues (Phe336 and Phe364) in the C-terminal domain, bidentate Fe(II) coordination, and deprotonation of the enol form. Recent crystal structures of the human HPD–NTBC complex further revealed extensive hydrophobic interactions within the catalytic cavity and ligand-induced rearrangements surrounding the C-terminal gating helix that stabilize inhibitor binding [7]. The resulting highly stable enzyme-inhibitor complex leads to near-irreversible enzyme inactivation with extremely slow recovery of HPD activity (Fig. 1d) [3,4,19–24].

Although NTBC dramatically improves survival in HT1, it neither corrects the underlying FAH deficiency [25] nor eliminates the risk of developing hepatocellular carcinoma [26–28]. NTBC withdrawal remains necessary in most gene- or cell-based therapies to create selective pressure for corrected hepatocytes, yet this exposes uncorrected cells to toxicity and facilitates malignant transformation. Because complete hepatocyte correction is unrealistic, NTBC remains indispensable while simultaneously limiting advanced therapeutic strategies [29,30].

To bypass the limitations of NTBC-dependent therapy, metabolic pathway reprogramming has emerged as an alternative strategy. Upstream inactivation of HPD via CRISPR-mediated disruption or *in utero* cytosine base editing can rescue the lethal HT1 phenotype in mice and pigs following NTBC withdrawal. However, this permanently introduces a secondary genetic lesion, producing an HT3-like metabolic state without correcting the underlying HT1-causing FAH mutation [31–34].

To overcome the drawbacks of both NTBC-dependent therapy and irreversible HPD inactivation, we aimed to engineer HPD variants capable of maintaining catalytic activity in the presence of NTBC. Using directed protein evolution (DPE) through the Knowledge Gaining Directed Evolution (KnowVolution) strategy, we generated NTBC-resistant HPD variants. KnowVolution integrates random mutagenesis, targeted saturation mutagenesis, computational modelling, and combinatorial recombination to efficiently identify and exploit positions contributing to a desired protein property [35]. Through iterative rounds of mutagenesis and high-throughput selection in our bacterial screening system[36], we identified eight substitutions that increased resistance to NTBC and related β-triketone inhibitors without compromising catalytic activity. The best-performing variant retained catalytic activity at NTBC concentrations far exceeding those tolerated by wild-type (wt) HPD *in vitro*.

By showing the feasibility of engineering NTBC-tolerant human HPD variants, this study tackles the main hurdle in the development of future gene- and cell-based therapies for HT1 that seek to maintain tyrosine catabolism. This strategy could eliminate the need for NTBC withdrawal and hereby reduces hepatotoxic risk. Such an advancement is particularly relevant given that residual FAH-deficient hepatocytes remain intrinsically prone to malignant transformation when exposed to accumulated toxic intermediates.

## 2. Methods

### 2.1 Materials

All chemicals were of analytical-reagent grade or higher quality and were purchased from Sigma-Aldrich Chemie GmbH (Schnelldorf, Germany), AppliChem (Darmstadt, Germany), or Carl Roth GmbH (Karlsruhe, Germany). All enzymes were purchased from New England Biolabs (Beverly, USA), Sigma Aldrich (Hamburg, Germany), or ThermoFisher Scientific (Waltham, MA, USA). NTBC was purchased from Yecuris (Tualatin, OR, USA). Oligonucleotides were purchased from Eurofins MWG Operon (Ebersberg, Germany) or ThermoFisher Scientific (Waltham, MA, USA) in salt-free form. A thermal cycler (Mastercycler gradient; Eppendorf, Hamburg, Germany) and thin-wall PCR tubes (Multi ultra-tubes; 0.2 mL; Carl Roth, Germany) were used in all PCRs. Plasmid extraction and PCR purification kits were purchased from Macherey-Nagel GmbH & Co. (Düren, Germany), Sigma Aldrich (Hamburg, Germany), and Qiagen (Hilden, Germany). DNA concentrations were quantified using a NanoDrop spectrophotometer (Thermo Scientific, Wilmington, USA). Microtiter plates for protein expression (96-wells; V-shaped) and for absorbance measurements (96-wells, flat-bottom) (Greiner Bio-One GmbH, Frickenhausen, Germany) were incubated in a Multitron II Infors shaker (Infors AG, Bottmingen, Switzerland) or Innova^TM^ 42R Incubator Shaker (Eppendorf, Hamburg, Germany). For detection of absorbance, Tecan Sunrise^®^ or Tecan Infinite^®^ M200 pro (Tecan Group Ltd, Männedorf, Switzerland) plate readers or Victor3 multilabel plate reader (Perkin Elmer, Waltham, MA, USA) were used.

### 2.2 Bacterial strains and plasmids

*Escherichia coli* (*E. coli*) DH5α (Stratagene, La Jolla, CA, USA) was used for DNA propagation, and *E. coli* C43 (DE3) (Lucigen Corporation, Middleton, WI, USA) for expression of recombinant enzymes and library screening. The expression vector pET42b(+) (Novagen, Darmstadt, Germany) carried a kanamycin resistance gene. As a negative control, empty pET42b(+) (EV) was transformed into *E. coli* [36].

### 2.3 Gene cloning and construction of expression vectors

cDNA of WT human HPD (NM_002150.2) was obtained from Geneart in a pMA-T-based cloning plasmid (Thermo Scientific), optimized for *E. coli* codon usage (see supplementary information 1). The HPD gene was subcloned as an NdeI-XhoI fragment into pET42b(+), transformed into *E. coli* DH5α and sequenced. Lysogeny Broth (LB) medium supplemented with kanamycin (50 μg/mL) was used for selection [36]. Sanger sequencing was performed by GATC Biotech AG (Cologne, Germany) using T7 FW (5’-TAA TAC GAC TCA CTA TAG GG -3’; Tm 53.2°C, GC 40%) and T7 RV (5’-GCT AGT TAT TGC TCA GCG GT - 3’; Tm 54.5°C, GC 47%). Alignments were performed using BioEdit 7.2.

### 2.4 Diversity Generation by KnowVolution

#### 2.4.1 Phase 1 - Construction of HPD epPCR library

Random mutagenesis was performed using epPCR [37] in tandem with phosphorothioate-based ligase-independent gene cloning[38] (PLICing; list of PTO-primers in supplementary data table S1). MnCl_2_ concentration (0–0.5mM) was optimized to achieve ∼50% active clones. PCR conditions for the HPD insert in the presence of Taq DNA polymerase to generate a mutant library were 94°C for 3 min; 30 cycles of 94°C 30 s, 48.5°C 30 s, 72°C 1 min; final elongation 72°C 10 min. Vector backbone amplification was performed in the presence of Phusion High-Fidelity DNA polymerase with the following cycling conditions: 98°C 3 min; 25 cycles of 98°C 30 s, 58.3°C 30 s, 72°C 2 min; final elongation 72°C 5 min. PCR products were verified by agarose gel electrophoresis [39], digested with DpnI (20 U, NEB), purified, quantified, and used in PLICing reactions with a 0.05 pmol/µL insert to 0.03 pmol/µL vector ratio. Products were transformed into chemically competent *E. coli* C43 (DE3) and plated on LB_KANA_. Religation and DpnI controls were included to monitor background colony formation.

#### 2.4.2 Phase 2 - Determination of beneficial substitutions by site-saturation mutagenesis

Nine positions (118, 137, 151, 153, 166, 177, 278, 302, and 307) identified from epPCR were subjected to SSM using 22c-trick primers [40] (NDT : VHG : TGG = 12:9:1, see supplementary data table S2 for primer sequences). A modified QuikChange two-step PCR protocol was applied with Phusion High-fidelity Polymerase[41]. PCR conditions for the first step: 98°C for 1 min; 3 cycles of 98°C 30 s, 60°C 30 s, 72°C 3 min. For the second step: 98°C 1 min; 18 cycles of 98°C 30 s, 60°C 30 s, 72°C 3 min; final elongation 72°C 8 min. After PCR, DpnI digestion removed template DNA. Libraries were transformed into *E. coli* C43 (DE3) and screened in 96-well plates using the previously described high-throughput assay [36]. An oversampling factor of three was applied to achieve ∼95% coverage of the theoretical variant space, requiring screening of ∼66 colonies per position [40].

#### 2.4.3 Phase 3 - Computational structural analyses

The crystal structure of human HPD was obtained from the Protein Data Bank (PDB ID: 3ISQ)[42]. Structural visualization, residue mapping, and figure preparation were performed using UCSF ChimeraX (version 1.11.1) [43]. The HPD-M8 structural model was generated by *in silico* mutagenesis of the HPD-WT structure using Rowan Scientific Boltz-2 based co-folding (https://www.rowansci.com accessed 2026) and subsequently visualized in ChimeraX. Experimentally identified substitutions from phase I of KnowVolution were mapped onto chain A of the HPD structure and classified according to their spatial localization relative to the catalytic cavity. Catalytic pocket residues were defined as residues located within 5 Å of either the Fe(II) cofactor or the substrate/inhibitor binding region. Surface residues were identified based on solvent exposure following visualization of the molecular surface in ChimeraX.

Protein stability effects of individual substitutions were assessed on the (i) Boltz-2 generated monomer, (ii) the crystal structure of human HPD-WT complexed with NTBC (8IM2) and (iii) a Boltz-2 generated model of HPD-WT homodimer complexed with TTC36 and Fe^2+^, using the DynaMut2 web server (https://biosig.lab.uq.edu.au/dynamut2/), which predicts mutation-induced changes in folding stability (ΔΔG, kcal/mol) from protein structural information [44]. Averaged positive ΔΔG values from all three models were interpreted as stabilizing and negative values as destabilizing. The predicted folding stability of HPD-WT and the final eight-substitution variant HPD-M8 was evaluated using IFUM (*in silico* evaluation of unfolding Free energy with Unfolded state ensemble Modeling), a deep-learning method that estimates absolute unfolding free energy (ΔG) by jointly considering sequence and structural representations of the folded and unfolded states [45] Predicted ΔG values were obtained for the complete wild-type and M8 sequences, and the stability difference (ΔΔG) was calculated as ΔG_M8_ − ΔG_WT_, with positive values indicating increased predicted stability of the variant. Potential changes in MHC class II peptide presentation were evaluated using NetMHCIIpan 4.1 BA mode (https://tools.iedb.org/mhcii/) across representative HLA-DR, HLA-DP, and HLA-DQ alleles [46]. For each variant, the number of predicted MHC-II binders was compared with that of HPD-WT. Substitutions were classified according to the net gain or loss of predicted binders.

Docking analyses of the native substrate HPP and the inhibitor NTBC were performed using the DiffDock/SMINA [47] pipeline implemented through the NeuroSnap platform (NeuroSnap Inc., 2022). Docking calculations were carried out for both HPD-WT and the engineered HPD-M8 variant. For each ligand-protein combination, multiple docking poses were generated and ranked according to docking rank and SMINA affinity score, where lower affinity scores indicate more favourable predicted binding interactions. Docking analyses were interpreted qualitatively to compare substrate and inhibitor accommodation patterns between HPD-WT and HPD-M8.

Structural interface analyses of the HPD biological assembly were performed using the PDBePISA server (v.1.52 10/20/2014) [48,49] to evaluate solvent accessibility surface area (ASA), buried surface area (BSA), and potential contributions of experimentally identified residues to the HPD dimer interface.

Tunnel and catalytic access pathway analyses were performed using CAVER Web[50] to investigate ligand accessibility of the buried catalytic cavity. HPD-WT and HPD-M8 structural models were co-folded with TTC36 [51] as molecular chaperone using Boltz-2 and analysed using both HPP, NTBC and HGA as ligands. The catalytic Fe(II) centre was used as the starting point for tunnel calculations. Resulting transport-energy profiles and energetic barriers along the catalytic access pathway were compared between HPD-WT and HPD-M8 to assess potential effects of the experimentally identified substitutions on substrate and inhibitor passage.

Evolutionary conservation analysis was performed using the ConSurf server [52]. Homologous sequences were identified with HMMER [53] (E-value ≤ 0.0001), filtered with CD-HIT [54] (≤ 95% sequence identity), and aligned using MAFFT [55]. A phylogenetic tree was generated using the Neighbor-Joining method with maximum-likelihood distance estimation [56], and site-specific conservation scores were calculated with Rate4Site [57]. Conservation scores of experimentally identified residues were compared with those of residues forming the catalytic centre and inhibitor-binding cavity of HPD.

Six positions (239, 241, 334, 363, 364, and 367) identified after molecular modelling (in the catalytic centre) were subjected to SSM using 22c-trick primers [40] (NDT : VHG : TGG = 12:9:1, see supplementary data table S3). A modified QuikChange two-step PCR protocol was applied [41]. PCR conditions for the first step: 98°C for 1 min; 3 cycles of 98°C 30 s, 60°C 30 s, 72°C 3 min. For the second step: 98°C 1 min; 18 cycles of 98°C 30 s, 60°C 30 s, 72°C 3 min; final elongation 72°C 8 min. After PCR, DpnI digestion removed template DNA. Libraries were transformed into *E. coli* C43 (DE3) and screened in 96-well plates using the previously described high-throughput assay [36].

#### 2.4.4 Phase 4 - Recombination of beneficial positions by multi-site-directed mutagenesis

Beneficial substitutions were combined to create triple (M3), quintuple (M5), and octuple (M8) HPD variants using a two-step PCR protocol with Phusion High-fidelity polymerase [41]. Primer mixes were designed for region 1 (positions 118, 138, 151, 153, 177) and region 2 (278, 302, 307) (see supplementary data table S4). PCR conditions for the first step: 98°C for 1 min; 3 cycles of 98°C 30 s, 60°C 30 s, 72°C 3 min. For the second step: 98°C 1 min; 18 cycles of 98°C 30 s, 60°C 30 s, 72°C 3 min; final elongation 72°C 8 min. After PCR, DpnI digestion removed template DNA. Libraries were transformed into *E. coli* C43 (DE3) and screened in 96-well plates using the previously described high-throughput assay [36]. PCR products were digested with DpnI and transformed into *E. coli* C43 (DE3) for expression.

### 2.5 High-throughput library screening

The basics of the HTS assay have been previously described [36]. Briefly, colonies were inoculated in 96-well plates with 100 µL LB + 1% glucose and incubated at 30°C, 900rpm, 70% RH for 16 h. Cultures were transferred to expression plates with 1mM isopropyl β-D-1-thiogalactopyranoside (IPTG) and 0.75 mg/mL Tyr, in the absence or presence of NTBC. After 24 h at 37°C, 900rpm, 70% RH, cultures were centrifuged, and OD_405_ was measured to quantify pyomelanin formation.

### 2.6 Confirmation re-screening of selected variants

Variants of interest were rescreened with increasing NTBC concentrations. Overnight cultures were diluted to OD_600_ 0.1, grown to OD_600_ 0.6– 0.8, induced with 1mM IPTG, and challenged with NTBC with four replicates of each variant per concentration in 96-well plates. Absorbance at OD_405_ was measured after 24 h [36].

### 2.7 Dose-response inhibition curves

HPD-WT and HPD-M8 were assayed against mesotrione (0 – 15µM), NTBC (0 – 20µM), sulcotrione (0 – 20µM), and tembotrione (0 – 40µM) with 4 replicates per concentration (n = 20). The normalized OD_405_ (OD_405-N_) was calculated according to OD_405-N_ = OD_405_ – OD_405-EV_ (Eq. 1) where OD_405-EV_ = measured OD_405_ of the EV. The activity percentage (Act. %) was calculated with GraphPad Prism 7 software according to the equation Act.% = (OD_405-N_/OD_405-M_) * 100 (Eq. 2), where OD_405-M_ is the mean of OD_405_ measured at 0µM (i.e. 100% activity)[36].

### 2.8 Whole-cell dose-response experiment with tyrosine

The HPD-M8 enzyme variant and the HPD-WT enzyme were challenged with different concentrations of tyrosine (0 – 10mM) with 4 replicates per concentration to assess substrate responsiveness. The same reaction was also set up, but with the addition of 2µM NTBC, i.e., the concentration where the HPD-WT is completely blocked and HPD-M8 is 100% active. The assay and measurements were performed as previously described [36].

### 2.9 Purification and sequencing of human HPD variants

Plasmids from overnight cultures were purified using the Nucleospin® Plasmid DNA Purification kit from Macherey-Nagel GmbH & Co. (Düren, Germany). The isolated plasmid DNA was quantified at 260nm and sequenced using T7 primers. Alignments were performed in BioEdit 7.2.

### 2.10 Statistical analysis

OD_405_ values were blank-corrected using EV+NTBC and normalized to WT or HPD-M8 without NTBC. Data are expressed as mean ± 95% CI. One-way ANOVA followed by Bonferroni post hoc test was applied (α = 0.05) using GraphPad Prism 11.0.1.

## 3. Results & Discussion

The aim of this study was to engineer human HPD variants with improved tolerance to NTBC and related β-triketones while preserving their native catalytic function.

Despite extensive structural analysis and multiple rational design attempts [2,3,5,6,10], the contribution of individual residues to NTBC binding or tolerance remains poorly understood. To address this, we applied the KnowVolution strategy, which integrates random mutagenesis, site-saturation mutagenesis, computational analysis, and recombination (Fig 2). This workflow efficiently identifies beneficial substitutions while reducing screening effort and often outperforms classical multi-round directed evolution campaigns [35].

**Fig. 2:**
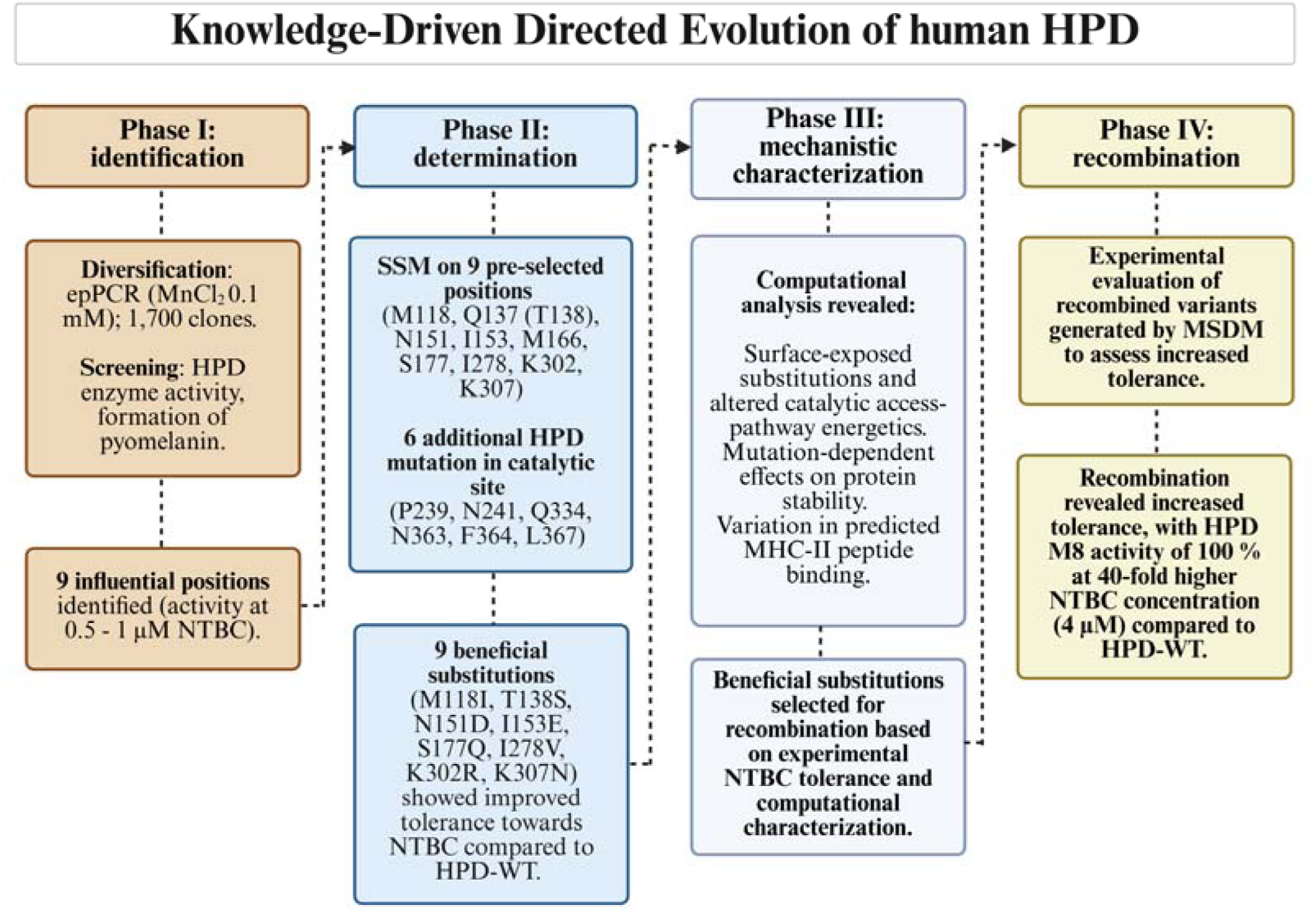
Schematic overview of the KnowVolution strategy applied to engineer NTBC-tolerant human HPD variants. In Phase I, random mutant libraries were generated by epPCR and screened using the previously described high-throughput assay [36] in the presence of NTBC, resulting in the identification of nine beneficial positions associated with increased inhibitor tolerance. In Phase II, site-saturation mutagenesis (SSM) was performed on the preselected positions, together with selected catalytic-site residues involved in NTBC accommodation, to identify beneficial amino acid substitutions while assessing tolerance of active-site modifications. In Phase III, computational structural analyses including PDBePISA, DiffDock/SMINA docking, DynaMut2, NetMHCIIpan, CAVER Web transport analyses, and ConSurf conservation analysis were used to investigate the mechanistic basis of NTBC tolerance. These analyses revealed that beneficial substitutions predominantly localize to solvent-exposed surface regions and indirectly modulate catalytic cavity accessibility and inhibitor transport energetics without disrupting the conserved catalytic machinery. In Phase IV, beneficial substitutions identified in Phases I and II were recombined by multi-site directed mutagenesis (MSDM), resulting in the HPD-M8 variant, which retained full catalytic activity at NTBC concentrations approximately 40-fold higher than those inhibiting HPD-WT. (Created in BioRender. Lequeue, S. (2026) https://BioRender.com/ad1l3q5).

### 3.1 Phase 1. Random mutagenesis identifies beneficial substitutions improving NTBC tolerance

Random mutant libraries were generated using 0.1mM MnCl_2_, low-fidelity DNA polymerase, and balanced dNTPs, and screened using a whole-cell colorimetric assay based on pyomelanin formation (Fig.3a – b) [36]. As pyomelanin production reflects HPD activity and is strongly inhibited by NTBC, pigment formation in its presence served as a proxy for inhibitor tolerance. Screening ∼1700 clones simultaneously without NTBC and at 0.25 – 0.50µM NTBC yielded nine variants with reproducible improvements in tolerance. A secondary rescreen across 0 – 1µM confirmed these observations. Sequencing identified several recurrent positions (i.e. 151 and 302), though with distinct amino-acid substitutions. The identified substitutions included M118I/N, Q137R, N151D, I153N, M166T, S177G, I278V, K302R, and K307N. Furthermore, comparison of variant activities in the absence and presence of NTBC revealed distinct effects on basal activity and inhibitor tolerance. I278V and N151D exhibited increased basal activity (∼115 – 120% of WT) and improved NTBC tolerance, retaining ∼71% and ∼45% of their respective uninhibited activities at 0.50µM NTBC, compared with ∼3% for WT. M118I also showed increased inhibitor tolerance despite reduced basal activity, whereas I153N, S177G, K302R, and K307N displayed increased basal activity and varying degrees of tolerance, particularly at 0.25µM NTBC. In contrast, M166T showed a modest increase in basal activity, but remained susceptible to inhibition at higher NTBC concentrations, while Q137R showed no apparent improvement in either property (Fig. 3c – f).

**Fig 3:**
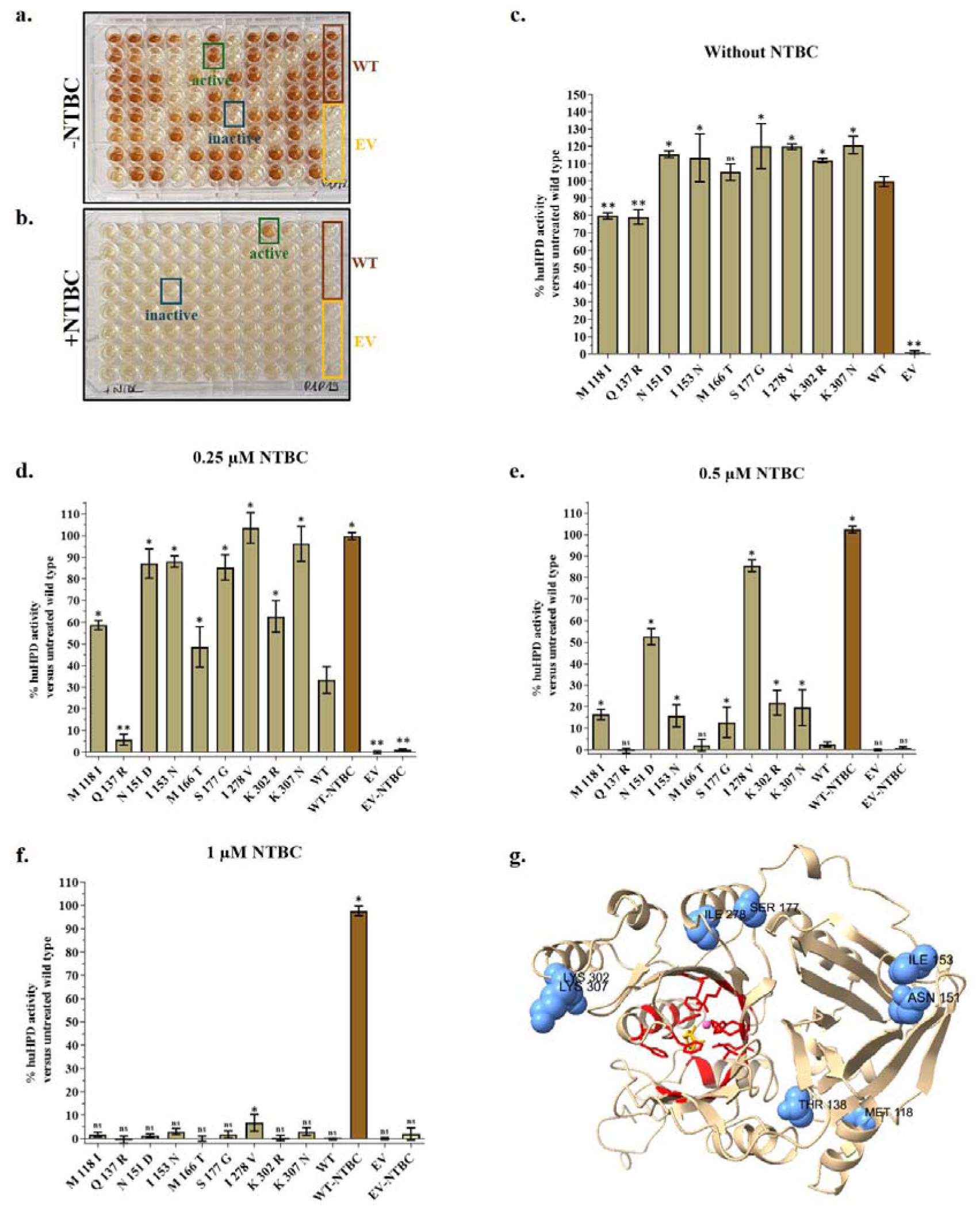
High-throughput screening of human HPD variant libraries in the (a) absence and (b) presence of 0.25µM NTBC resulting in inactive and active (improved) variants. Activity and tolerance rescreening of newly identified nine human HPD enzyme single variants in the presence of (c) 0 – (d) 0.25 – (e) 0.5 – (f) 1µM NTBC. Four replicates of each variant were used in the rescreening assay and challenged with the different concentrations of NTBC. Several variants still retained approximately 20% activity at 0.5µM NTBC, while the HPD-WT is almost completely blocked. The data are represented as mean ± 95% CI. Statistical significance was determined by a one-way ANOVA with Bonferroni correction.* Significantly higher activity than HPD-WT (p < 0.05), ** significantly lower activity than HPD-WT (p < 0.05). Abbreviations: WT: wild-type human HPD enzyme exposed to NTBC; WT-NTBC: wild-type human HPD enzyme not exposed to NTBC, considered as 100% active; EV: empty vector control exposed to NTBC; EV-NTBC: empty vector control not exposed to NTBC. (g) Structural mapping of residues identified in Phase I on chain A of the human HPD monomer. The catalytic pocket (defined as residues located within 5 Å of both the cofactor and substrate) is shown in red and identified residues in blue. The substrate (HPP) is shown in yellow, and the non-heme Fe^2+^ cofactor is shown in pink. None of the residues were located within the catalytic pocket.

Overall, I278V and N151D exhibited the most pronounced combined improvements in activity and NTBC tolerance, with I278V retaining 6.8 ± 2.2% residual activity at 1µM NTBC, although all variants were strongly inhibited at 1µM NTBC (Fig. 3c – f). To gain structural and mechanistic insight, the identified positions were mapped onto the 3D monomeric structure of HPD. All identified residues were solvent-exposed and spatially separated from the catalytic pocket. Notably, none of the identified positions were located within the catalytic pocket (Fig. 3g).

### 3.2 Phase 2. Site-saturation mutagenesis refines beneficial NTBC-tolerant substitutions

Targeted saturation mutagenesis was performed at the nine positions identified in phase 1. Substitutions at positions 153 (I153E) and 307 (K307P) generated the largest increases in NTBC tolerance, and T138S emerged as an additional beneficial substitution. Positions 118, 151, 166, 177, or 302 did not yield enhanced tolerance upon systematic substitution (Fig. 4). To ensure ∼95% coverage of the variant space, at least 66 clones per position were required [40]. In practice, 88 SSM clones per position were screened across 0–2µM NTBC. This confirmed the phase 1 findings and narrowed the key positions and substitutions for further combinatorial optimization.

**Fig. 4:**
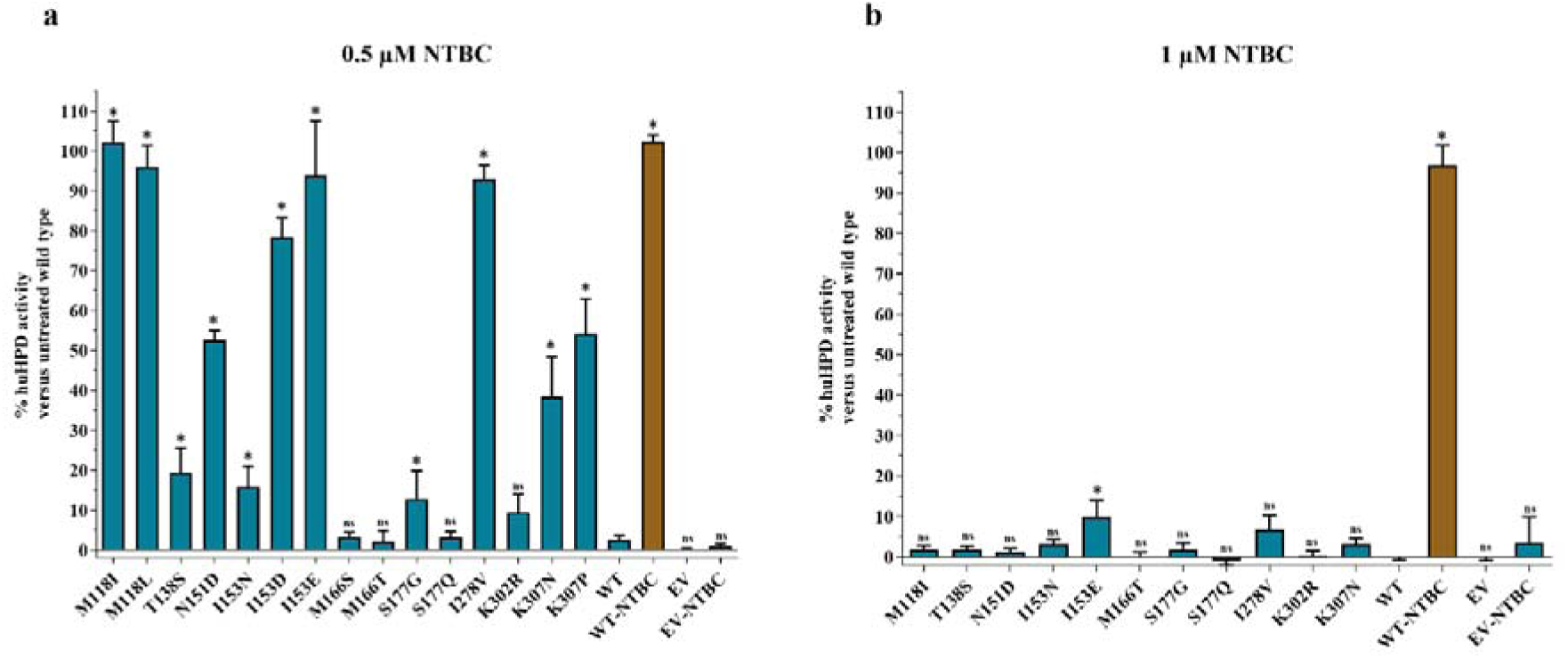
Activity and tolerance after SSM of the nine human HPD enzyme variants in the presence of (a) 0.5 and (b) 1µM NTBC. Four replicates of each variant were used in the rescreening assay and challenged with the different concentrations of NTBC. The data are represented as mean ± 95% CI. Statistical significance was determined by a one-way ANOVA with Bonferroni correction.* Significantly higher activity than HPD-WT (p < 0.05). Abbreviations: WT: wild-type human HPD enzyme exposed to NTBC; WT-NTBC: wild-type human HPD enzyme not exposed to NTBC, considered as 100% active; EV: empty vector control exposed to NTBC; EV-NTBC: empty vector control not exposed to NTBC.

**Fig. 5:**
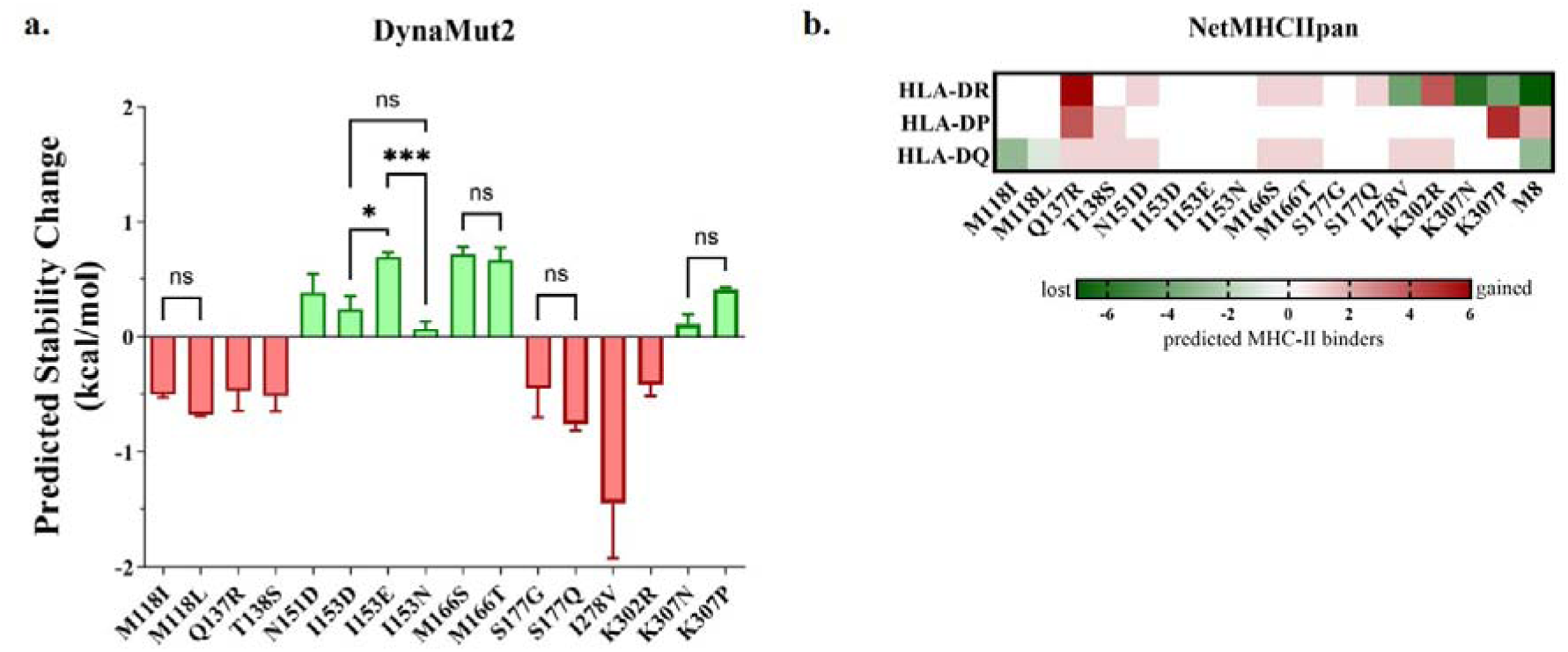
*In silico* analysis of the effects of identified HPD substitutions on protein stability and predicted MHC-II binding. (a) Changes in protein stability predicted using DynaMut2. Positive ΔΔG values indicate stabilizing substitutions, whereas negative values indicate destabilizing substitutions. Statistical significance between substitutions at the same amino-acid position is indicated as ns, not significant; *p < 0.05; and ***p < 0.001. (b) Changes in the number of predicted MHC-II binders determined using NetMHCIIpan for HLA-DR, HLA-DP, and HLA-DQ alleles. Values represent the net change in predicted binders relative to HPD-WT, with green indicating a loss and red indicating a gain of predicted MHC-II binders. M8 represents the lead variant containing eight accumulated substitutions.

### 3.3 Phase 3. Computational evaluation of individual substitutions reveals differential effects on protein stability and MHC-II binding

Individual substitutions identified in the preceding two phases were evaluated computationally with respect to their predicted effects on protein stability and MHC-II peptide presentation to guide subsequent recombination. DynaMut2 analysis predicted that N151D, I153D/E/N, M166S/T, and K307N/P were to be stabilizing or near-neutral, with the largest favourable stability changes observed for M166S (+0.72 ± 0.06 kcal/mol), I153E (+0.70 ± 0.04 kcal/mol), and M166T (+0.67 ± 0.11 kcal/mol). In contrast, M118I/L, Q137R, T138S, S177G/Q, I278V, and K302R were predicted to be destabilizing, with I278V showing the largest predicted stability penalty (−1.45 ± 0.48 kcal/mol) (Fig.5a; Table S5). NetMHCIIpan analysis further revealed mutation-specific changes in the predicted HLA-II binder profile. I153D/E/N and S177G did not alter the total number of predicted binders, whereas M118I, I278V, and K307N reduced the predicted binder burden by 3, 3, and 6 binders, respectively. Conversely, Q137R and K302R produced the largest increases, with net gains of 11 and 5 predicted binders across the HLA-DR, -DP, and -DQ panels (Fig.5b; Fig.S1).

### 3.4 Phase 4. Recombination of beneficial substitutions yields highly NTBC-tolerant HPD variants

Based on the knowledge gained within the previous three phases of KnowVolution, multi-site directed mutagenesis (MSDM) was performed to combine all beneficial substitutions into a single HPD variant with high NTBC tolerance. The positions were grouped into two regions: R1 (positions 118, 138, 151, 153, 177 – M5) and R2 (positions 278, 302, 307 – M3). Using HPD-M3 as a template, all R1 positions were combined to generate the final variant HPD-M8, containing all beneficial changes. The resulting clones (M5, M3, and M8) were assayed across 1–6µM NTBC. HPD-WT was fully inhibited at 1µM NTBC. M5 retained 100% activity at 1–2µM and 30–40% at 3µM, whereas M3 showed 75% activity at 1µM and was fully blocked at 2–3µM. HPD-M8 retained full activity at 4µM NTBC, ∼50% activity at 5µM, and ∼5% at 6µM, corresponding to an approximate 40-fold improvement relative to WT (Fig. 6).

**Fig. 6:**
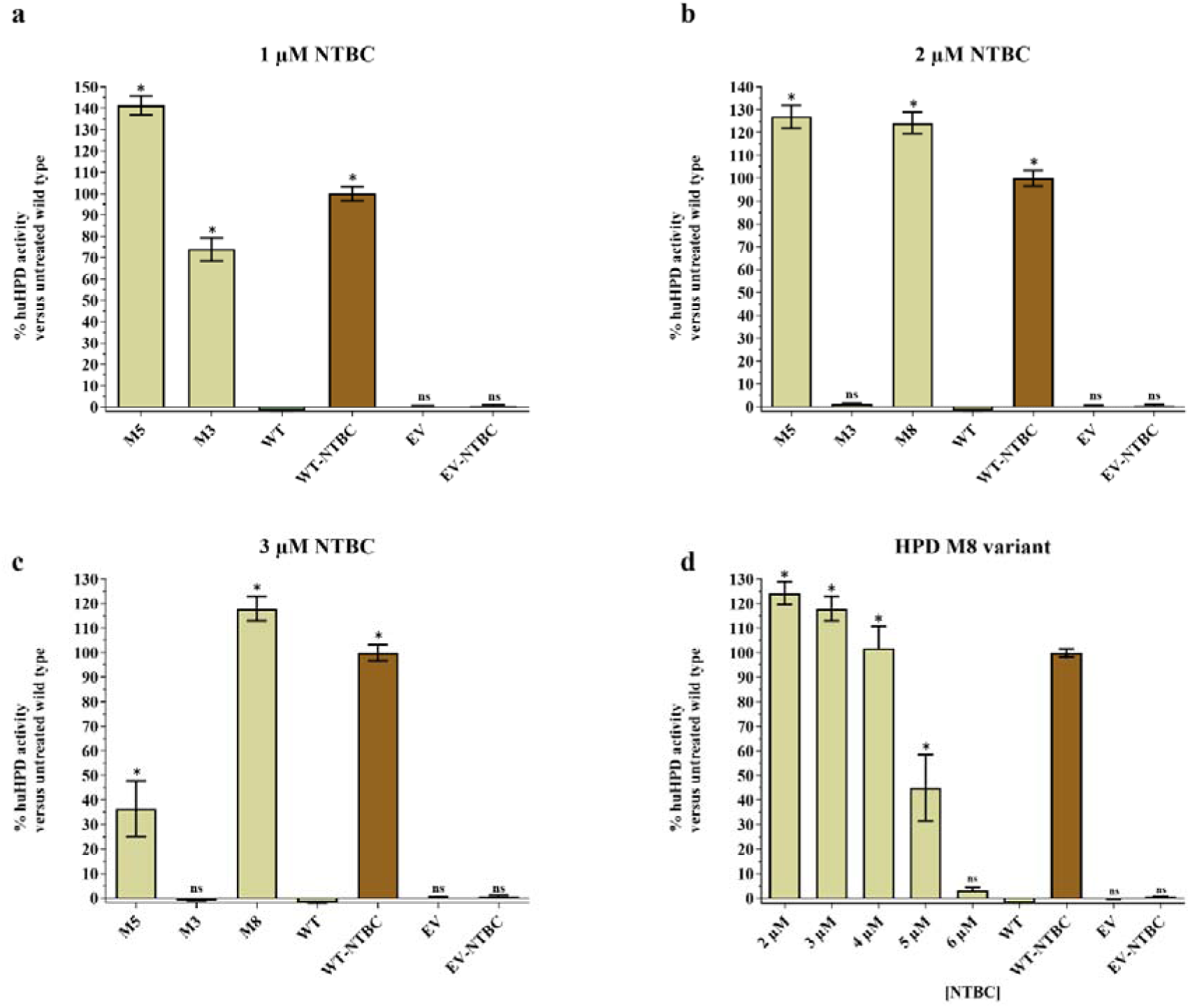
Rescreening results after performing MSDM where all identified positions and amino acid substitutions are recombined in multiple HPD variants, i.e. M5 (M118I, T138S, N151D, I153E, S177Q), M3 (I278V, K302R, K307N), and M8 (all positions and substitutions). Four replicates of each variant were challenged at different concentrations of NTBC: (a) 1µM, (b) 2µM, (c) 3µM. (d) The HPD-M8 variant is challenged at different concentrations of NTBC, while the WT version is completely blocked. The data are represented as mean ± 95% CI. Statistical significance was determined by a one-way ANOVA with Bonferroni correction.* Significantly higher activity than HPD-WT (p < 0.05). Abbreviations: WT: wild-type human HPD enzyme exposed to NTBC; WT-NTBC: wild-type human HPD enzyme not exposed to NTBC, considered as 100% active; EV: empty vector control exposed to NTBC; EV-NTBC: empty vector control not exposed to NTBC.

### 3.5 Computational analyses reveal distal modulation of catalytic cavity accessibility and no overall increased MHC-II binder burden

Computational analyses were performed to investigate how surface substitutions improve NTBC tolerance without compromising catalytic activity. Human HPD functions as a homodimer (PDB ID: 3ISQ)[42] and contains a highly conserved catalytic centre coordinating Fe(II) via His183, His266, and Glu349. Molecular docking confirmed that both HPP and NTBC interact within the same Fe(II) centred pocket (Fig. 1c-d). Consistent with previous structural studies, HPP formed interactions with residues including N241, G360, and Q334, whereas NTBC additionally interacted with residues lining the hydrophobic catalytic cavity, including P239, N241, Q334, N363, F364, and L367. Whole-protein docking analyses using DiffDock/SMINA showed that HPP retained favourable low-rank docking poses in both HPD-WT and HPD-M8, consistent with preserved substrate accommodation. NTBC docking produced a broader and more dispersed distribution of docking poses, particularly in HPD-M8 (Fig. 7), indicating greater variability in predicted inhibitor-binding conformations. However, as these docking analyses do not account for ligand transport, catalytic gating, or slow-binding kinetics, they were interpreted as supportive rather than definitive mechanistic evidence.

**Fig. 7:**
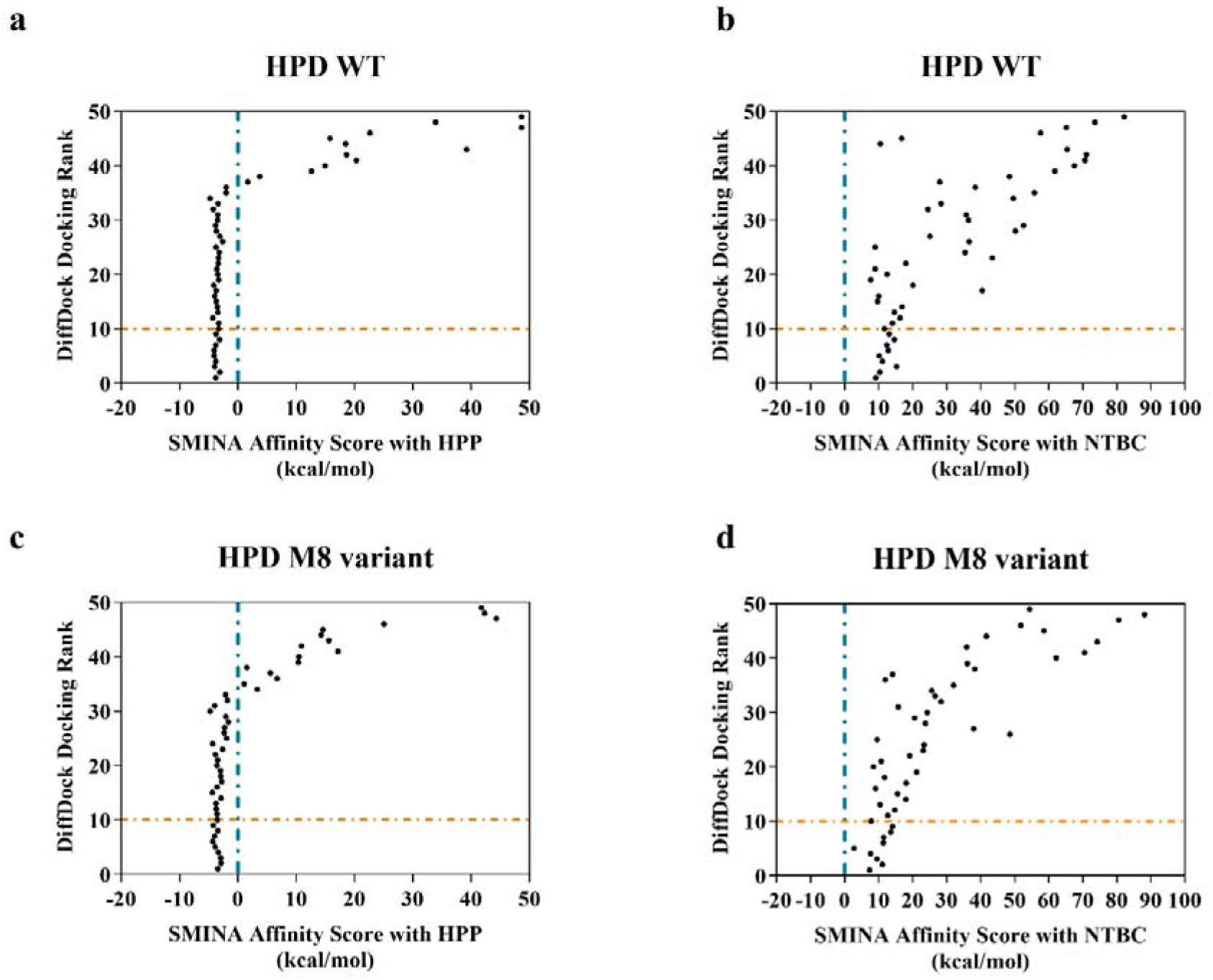
DiffDock/SMINA whole-protein docking analyses of the native substrate 4-hydroxyphenylpyruvate (HPP) and the inhibitor nitisinone (NTBC) in human HPD-WT and the engineered HPD-M8 variant. (a) Docking distribution of HPP in HPD-WT. (b) Docking distribution of NTBC in HPD-WT. (c) Docking distribution of HPP in HPD-M8. (d) Docking distribution of NTBC in HPD-M8. Each point represents an individual docking pose generated by the DiffDock/SMINA pipeline. Docking poses are plotted according to DiffDock docking rank and corresponding SMINA affinity score, where lower docking ranks and lower affinity scores indicate more favourable predicted ligand accommodation. HPP retained favourable low-rank docking poses in both HPD-WT and HPD-M8, consistent with preservation of productive substrate accommodation. In contrast, NTBC docking profiles appeared more dispersed and less consistently favourable, particularly in HPD-M8, indicating increased sensitivity of inhibitor accommodation to the engineered substitutions relative to substrate positioning. Because these docking analyses do not account for ligand transport, catalytic gating, or slow-binding kinetics, they were interpreted as supportive rather than definitive mechanistic evidence.

Structural mapping revealed that beneficial substitutions cluster on the protein surface and remain spatially separated from the catalytic pocket (Fig 3g). None of the substitutions directly contacted either the substrate or inhibitor, indicating that NTBC tolerance is not mediated through direct perturbation of catalytic-site interactions. Most identified positions localized to flexible loop regions (Met118, Asn151, Ile153, Met166, Ser177, and Lys307), α-helices (Ile278 and Lys302), or β-sheet-associated surface regions (Gln137 and Thr138), suggesting a potential role in modulating protein dynamics or catalytic cavity accessibility. PDBePISA analysis demonstrated that all beneficial substitutions are solvent-exposed and do not contribute to the dimer interface (BSA ∼ 0), suggesting that NTBC tolerance is unlikely to arise from direct perturbation of inhibitor-binding residues or altered dimerization.

To further investigate whether distal surface substitutions indirectly influence ligand accessibility, transport analyses were performed using CAVER Web and its docking module CaverDock (Fig. 8). Not surprisingly, the ligand transport tunnel, connecting the surface to the catalytic pocket, was significantly altered. Compared to HPD-WT, the bottleneck radius of HPD-M8’s ligand transport tunnel increased from 1.4 Å to 1.5 Å corresponding to an increase in throughput from 0.46 to 0.55 (Fig. 8a-g). The entrance region of the tunnel was energetically favourable in both proteins. Within the first 5 Å from the protein surface, HPD-WT and HPD-M8 showed comparable binding energy values, suggesting that the surface-proximal tunnel opening does not account for the major difference between the two systems (Fig. 8h-i). The main difference occurred in the central part of the tunnel. In HPD-WT, NTBC encountered a pronounced transport barrier, with binding energy reaching 31.5 kcal/mol at 11.39 Å from the tunnel entrance. This high-energy region extended across a broad central segment of the pathway, with binding energy remaining above 20 kcal/mol between approximately 9.69 Å and 15.46 Å (Fig. 8h-i). This central barrier coincided with tunnel narrowing, and the narrowest point of the HPD-WT trajectory was observed at 14.48 Å, where the tunnel radius decreased to 1.4 Å (Fig. 8g). In contrast, HPD-M8 displayed a substantially lower maximum binding energy of 9.9 kcal/mol at 11.83 Å from the tunnel entrance. Although the mutant tunnel also contained constricted regions, its energy profile did not reproduce the high central barrier observed in HPD-WT (Fig. 8h-i). The narrowest point in HPD-M8 occurred earlier along the transport pathway, at 7.91 Å, with a minimum radius of 1.5 Å (Fig. 8g). In addition, HPD-M8 showed a later moderate-energy region around 18 Å to 19.5 Å, suggesting redistribution of transport constraints rather than simple occlusion of the tunnel (Fig. 8h-i).

**Fig. 8:**
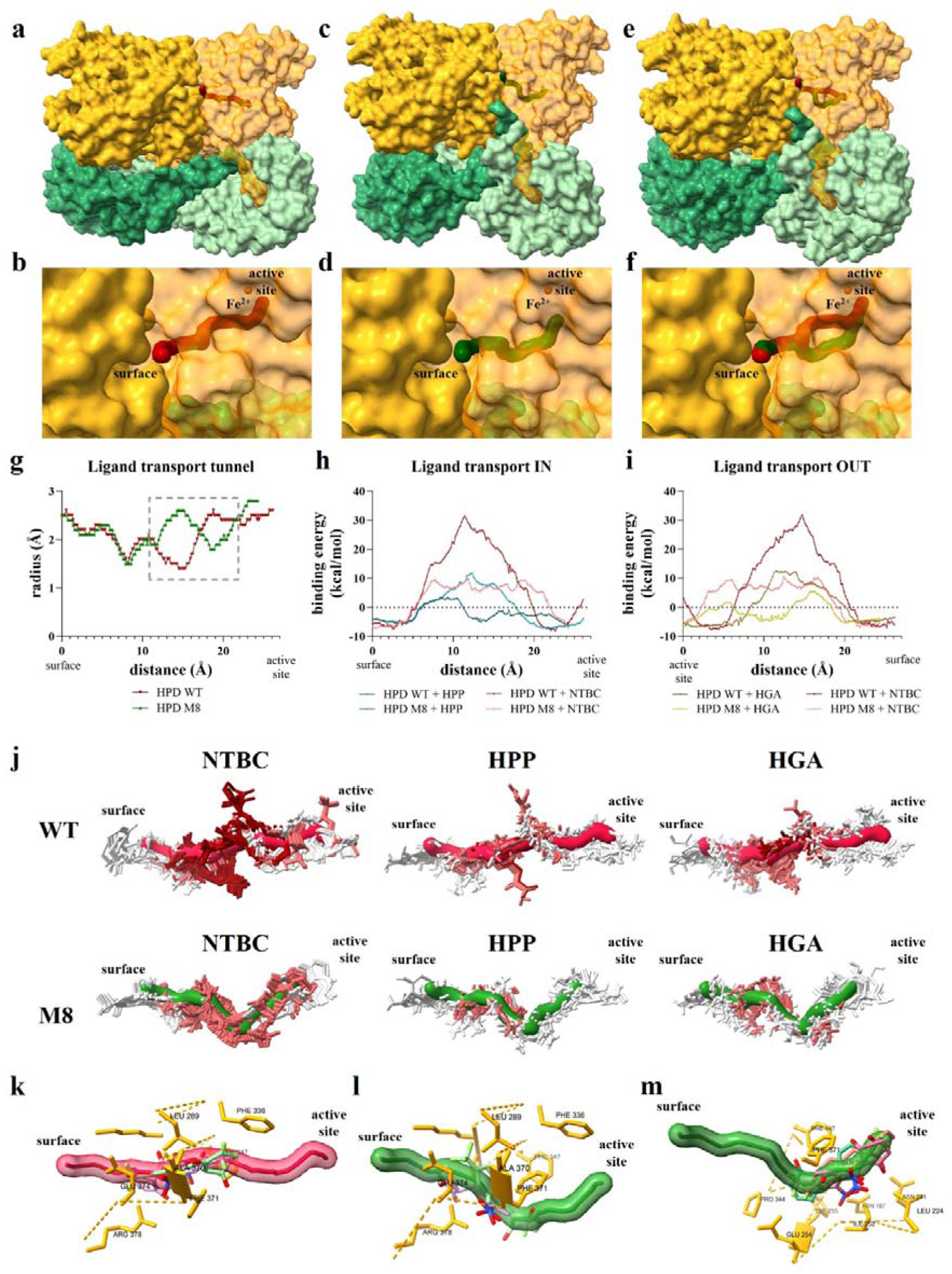
CAVER Web ligand transport tunnel analyses using Boltz-2 prediction models of human wild-type HPD homodimer or HPD-M8 (gold) co-folded with the molecular chaperone TTC36 (green). Visualization of the ligand transport tunnel connecting the surface and the buried active site of (a, b) HPD-WT, (c, d) HPD-M8 and (e, f) their superposition (g). Tunnel geometry significantly differs between HPD-WT and M8 from 11 Å to 22 Å distal from the surface (h-i). CaverDock analyses and (j) pose visualization of NTBC, HPP, and HGA transport through the predicted access tunnel of HPD-WT and HPD-M8. Transport-energy profiles were calculated along this catalytic access and egress pathway. Transport profiles of HPP and HGA were predicted to be positively enhanced in HPD-M8 *versus* HPD-WT. (k) HPD-WT displayed a pronounced central NTBC transport barrier, with binding energy reaching 31.5 kcal/mol at 11.39 Å and remained elevated across the central tunnel segment. This region coincided with tunnel narrowing and a major NTBC pose inversion between approximately 11.39 and 11.59 Å. (l) In contrast, HPD-M8 showed a shorter transport trajectory and a lower maximum binding energy of 9.9 kcal/mol at 11.83 Å. (m) The M8 mutant profile lacked the dominant WT-like central barrier, but showed redistributed moderate-energy regions, including a later constraint around 18 to 19.5 Å. Representative NTBC, HPP and HGA poses illustrate mutation-dependent differences in ligand orientation and accommodation along the tunnel pathway.

In addition, the maximal energy barriers for HPP decreased in HPD-M8 from 11.8 kcal/mol to 3.4 kcal/mol compared to HPD-WT and for HGA from 12.5 kcal/mol to 5.4 kcal/mol while binding energies near the surface and active site remained relatively unchanged (Fig. 8h-i). This suggests that the mutant tunnel remains competent for normal substrate entry and product passage, and that the M8 mutations are unlikely to impair the native catalytic cycle at the level of ligand transport.

Analysis of the NTBC poses along the tunnel further supported distinct ligand-accommodation patterns in the two proteins. In HPD-WT, the major NTBC reorientation occurred within the high-energy central barrier, between approximately 11.39 Å and 11.59 Å from the tunnel entrance. Across this transition, NTBC underwent an almost complete inversion of its aromatic-to-dione molecular axis, indicating that passage through this region requires a substantial pose rearrangement (Fig. 8k). This inversion was accompanied by changes in the relative positioning of the NTBC dione, nitro, trifluoromethyl and aromatic groups with respect to tunnel-lining residues, including Leu289, Phe336, Lys338, Pro339, Phe347, Ala370, Phe371, Glu374, and Arg378 (Fig.8k). HPD-M8 also exhibited NTBC reorientation near the central part of the tunnel, between approximately 11.83 Å and 11.99 Å (Fig. 8l). However, this transition occurred at a markedly lower energy than in HPD-WT. Additional pose rearrangements were observed in the later tunnel segment, particularly between 18.9 Å and 19.5 Å, where NTBC passed through a secondary moderate-energy region and shifted poses two times. These later rearrangements involved residues located deeper in the transport pathway, including Asn187, Leu224, Asn241, Ile252, Glu254, Tyr255, Gln265, Pro344, Phe347 and Phe371 (Fig. 8m). The M8 substitutions were not predicted to contact NTBC directly along the sampled transport trajectory. Instead, the observed differences are most consistent with an indirect structural effect, in which the M8 variant remodels the tunnel geometry and alters the sequence of NTBC poses adopted during transport. As such, HPD-M8 appears to shorten and reshape the access pathway, reduce the dominant WT-like central energy barrier and redistribute ligand-accommodation constraints toward additional tunnel regions. These findings suggest that the M8 substitutions alter NTBC access and positioning through long-range effects on tunnel architecture rather than through direct mutation/inhibitor contacts. Such changes may reduce the efficiency of NTBC accommodation and contribute to the observed inhibitor tolerance while maintaining substrate accessibility.

Evolutionary conservation analysis using ConSurf further demonstrated that most experimentally identified positions are weakly conserved among HPD homologues, with Lys302 representing the main exception. In contrast, residues forming the catalytic centre and inhibitor-binding cavity remained highly conserved. These observations support a model in which NTBC tolerance arises through substitutions at evolutionarily permissive surface regions that modulate structural dynamics without disrupting the conserved catalytic machinery.

The folding stability of the final HPD-M8 variant was further evaluated using IFUM [45]. HPD-WT showed a predicted unfolding free energy (ΔG) of 28.144 kcal/mol, whereas HPD-M8 displayed a higher predicted ΔG of 30.153 kcal/mol, corresponding to a ΔΔG of +2.009 kcal/mol. The higher IFUM-predicted unfolding free energy of HPD-M8 compared with wild-type HPD suggests that accumulation of the eight beneficial substitutions did not compromise overall protein stability and may instead confer a modest stabilizing effect. This is particularly relevant, because DynaMut2 predicted both stabilizing and destabilizing effects for the individual substitutions, indicating that their combined effect in HPD-M8 cannot be inferred solely from single-mutation predictions.

The predicted MHC-II presentation profile was additionally assessed for the final lead variant HPD-M8, which contained eight accumulated substitutions (Fig.5b) [46]. NetMHCIIpan predicted a net reduction of eight MHC-II binders relative to HPD-WT, comprising losses of seven HLA-DR and three HLA-DQ binders that were partially offset by the gain of two HLA-DP binders. The strongest reductions were associated with HLA-DRB112:01, HLA-DQA101:02/DQB106:02, and HLA-DRB501:01, whereas only limited allele-specific increases were observed (Fig.S1). Consequently, accumulation of the eight beneficial substitutions did not increase the overall predicted MHC-II binder burden, indicating that the enhanced functional properties of HPD-M8 were retained without a concomitant increase in predicted HLA-II peptide presentation across the investigated allele panel.

To determine whether direct perturbation of NTBC-contacting residues could similarly improve inhibitor tolerance, saturation mutagenesis was additionally performed on active-site residues involved in NTBC accommodation (Pro239, Asn241, Gln334, Asn363, Phe364, and Leu367). However, most substitutions resulted in inactive enzymes or variants indistinguishable from HPD-WT, indicating that direct modification of the catalytic pocket is poorly tolerated. Similar observations have previously been reported for homologous HPD enzymes, where active-site substitutions disrupt substrate positioning and reduce catalytic activity [5,58,59]. Collectively, these findings indicate that NTBC tolerance in HPD is mediated through long-range modulation of catalytic cavity accessibility rather than direct disruption of inhibitor-binding interactions.

### 3.6 Dose-response analysis with triketones

Inhibition of HPD-WT and HPD-M8 by triketones was evaluated in IPTG-induced *E. coli* expressing the enzyme [36]. Although NTBC is the only clinically relevant inhibitor, mesotrione, sulcotrione, and tembotrione were included to assess the broader tolerance of HPD-M8 [4]. Inhibitors were applied at 0–15µM (mesotrione), 0–20µM (NTBC, sulcotrione), and 0–40µM (tembotrione), with four replicates per concentration (Fig. 9a – d). Pyomelanin formation, reflecting HPD activity, decreased with increasing inhibitor concentration; bacterial cells were centrifuged prior to colorimetric quantification to prevent interference. IC_50_ values were determined using four-parameter logistic fits. HPD-WT IC_50_s were 0.363 ± 0.013µM (mesotrione), 0.195 ± 0.003µM (NTBC), 0.164 ± 0.015µM (sulcotrione), and 0.611 ± 0.015µM (tembotrione). HPD-M8 IC_50_s increased substantially to 5.103 ± 0.092µM, 3.933 ± 0.499µM, 3.249 ± 0.397µM, and 5.504 ± 0.174µM (Fig. 9e), respectively, corresponding to fold changes of 14–20 for mesotrione, NTBC, and sulcotrione, and ∼9 for tembotrione. Thus, HPD-M8 is broadly β-triketone tolerant, not NTBC-specific.

**Fig. 9:**
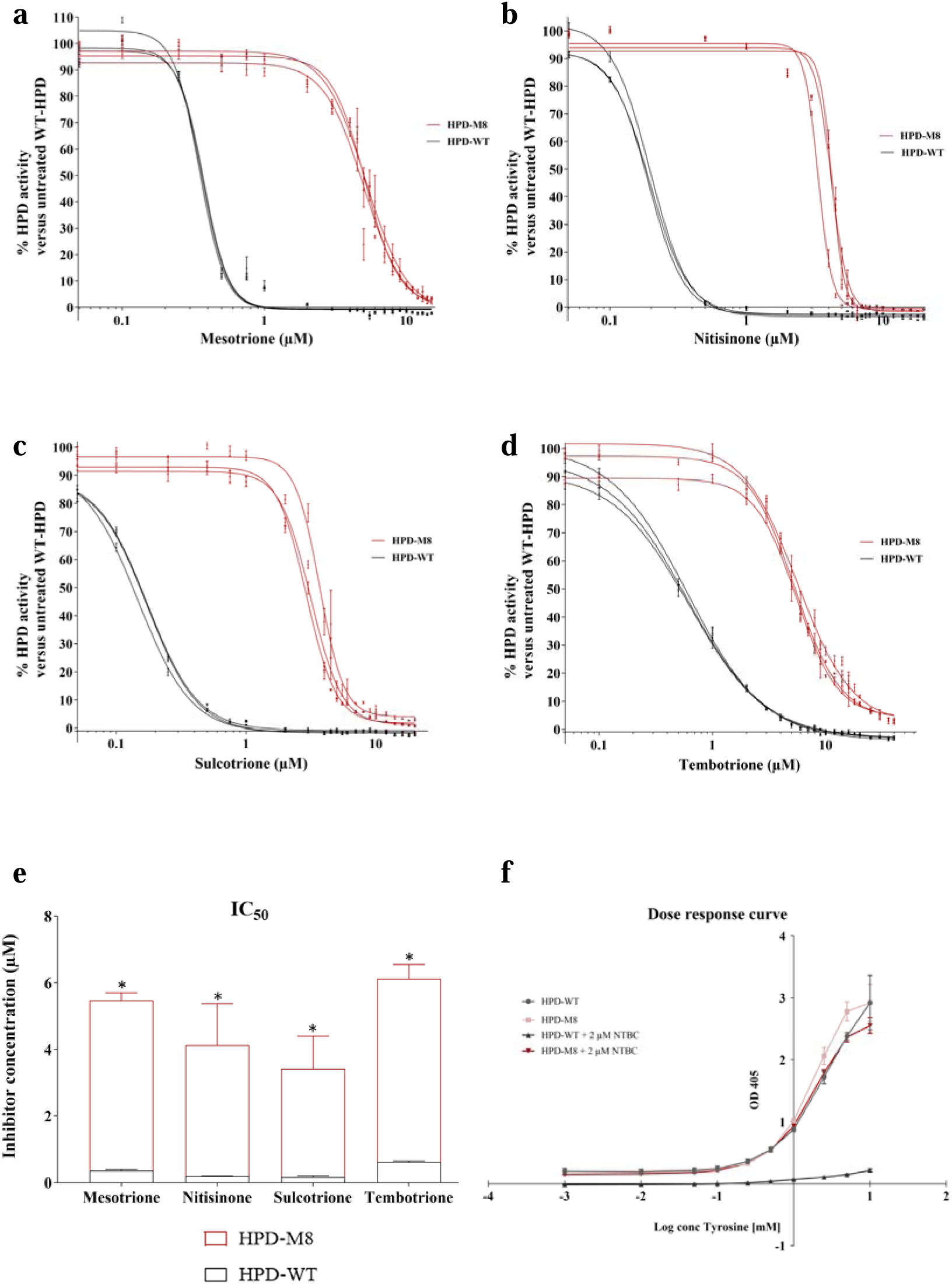
Dose-response inhibition curves of the prototypical β-triketone inhibitors (a) mesotrione (0 – 15µM), (b) nitisinone (0 – 20µM), (c) sulcotrione (0 – 20µM), and (d) tembotrione (0 – 40µM) for HPD-WT and HPD-M8. (e) The calculated IC_50_ values for each of the HPD inhibitors (n = 4). The data are represented as mean ± 95% CI. Statistical significance was determined by a two-way ANOVA with Holm-Sidak correction (P < 0.05). (f) Comparative activity analysis of HPD-WT and the engineered HPD-M8 variant at increasing tyrosine concentrations in the presence of a fixed NTBC concentration (2µM). Residual HPD activity was assessed using the whole-cell pyomelanin-based bio-assay [36] to evaluate whether the engineered substitutions altered the response of HPD-M8 to increasing substrate availability under inhibitory conditions. While HPD-WT activity remained strongly inhibited by NTBC, HPD-M8 retained substantially higher activity across increasing tyrosine concentrations, consistent with enhanced NTBC tolerance while preserving catalytic function. Data are represented as mean ± 95% CI from four biological replicates.

### 3.7 Substrate responsiveness

To assess whether HPD-M8 exhibited altered substrate responsiveness, HPD-WT and M8 were exposed to 0 – 10mM tyrosine. Since tyrosine availability determines HPP formation upstream, deviations could indicate altered substrate affinity. Both WT and M8 produced comparable pyomelanin levels across all conditions, in the presence or absence of 2µM NTBC (Fig. 9f). These findings indicate that HPD-M8 retains WT-like responsiveness to tyrosine and does not show obvious changes consistent with a modified Km.

### 3.8 Limitations and methodological considerations

A key limitation is the lack of direct kinetic characterization (Km, kcat, kcat/Km). Subtle effects on catalytic efficiency therefore cannot yet be excluded, especially given the complex multi-step inhibition of HPD by NTBC [1,4]. Although IC_50_ measurements provide comparative insights, more detailed kinetic analysis, *via* coupled-enzyme assays [10,58,60,61], oxygen consumption, HPLC methods [2,6], or composite kinetic metrics such as ON/OFF rate ratios, are required to definitively confirm that catalytic properties were unaltered [62].

The use of a bacterial evolution platform is another consideration. Although prokaryotic and yeast platforms are standard in directed evolution due to rapid growth and efficiency [63–66], they lack the native mammalian folding, degradation, localization machinery, and post-translational modifications (PTMs) [63,66]. Human HPD contains several PTMs, three phosphoserines (211, 226, 250), one N6-succinyllysine (132), and an N-acetyl-threonine (2) [1,4], none of which occur at the mutated positions, suggesting minimal risk of PTM interference. Structural differences between bacterial and mammalian HPD homologs, including tetrameric assembly in *P. fluorescens versus* dimeric organization in mammals [4,23,67–69], may also contribute to system-specific behaviour. Nonetheless, bacterial platforms remain effective for enzyme engineering when coupled to a robust, phenotype-linked assay [63–67], as demonstrated by the successful evolution of HPD-M8. Another limitation is that the NetMHCIIpan analysis only predicts peptide–MHC-II binding and therefore does not account for antigen processing, peptide abundance, T-cell receptor recognition or other factors that ultimately determine immunogenicity *in vivo*. Finally, the CAVER/CaverDock data should be interpreted as a predicted transport model, not as direct evidence of ligand diffusion dynamics, in support of a structural explanation for the reduced sensitivity of HPD-M8 to triketone inhibitors.

## 4. Conclusion

In this study, human HPD was successfully engineered using the KnowVolution strategy, yielding variants with markedly increased tolerance to NTBC and other β-triketone inhibitors. The final HPD-M8 variant retained full *in vitro* catalytic activity at NTBC concentrations at least 40-fold higher than those that completely inhibit HPD-WT. These results demonstrate that directed evolution can be applied to human metabolic enzymes to overcome clinically relevant inhibitor sensitivity, thereby opening new possibilities for designing gene-and cell-based therapeutic strategies for monogenic metabolic diseases.

## 5. CRediT authorship contribution statement

Conceptualization, J.N. and J.D.K.; Methodology, J.N., A.M., J.D.K.; Validation, J.N.; Formal Analysis, J.N. and J.D.K.; Investigation, J.N., A.M., J.D.K; Resources, T.V., U.S., and J.D.K.; Data Curation, J.N. and J.D.K.; Writing – Original Draft Preparation, J.N. and J.D.K.; Writing – Review and Editing, J.N., A.M., S.L., L.D., N.S.S., G.G.D, K.T., T.V., U.S. and J.D.K; Visualization, J.N., S.L., and J.D.K.; Supervision, T.V., U.S., and J.D.K.; Project administration, J.N.; Funding Acquisition, T.V. and J.D.K. All authors have read and approved the final manuscript.

## 6. Acknowledgements

The authors thank Dr. Gaurao V. Dhoke for his valuable assistance with the ConSurf evolutionary conservation analysis and helpful discussions regarding the interpretation of the results.

## 8. Declaration of generative AI and AI-assisted technologies in the writing process

During the preparation of this work, the authors used ChatGPT in order to improve the clarity and readability of certain sentences by refining grammar and restructuring complex phrases. After using this tool, the authors reviewed and edited the content as needed and take full responsibility for the content of the published article.

## 9. Additional Information

### Data Availability Statement

Tunnel analyses were performed using CAVER Web. Full CAVER Web outputs are available under the following job IDs: li7kgb (HPD-WT) and ri65f3 (HPD-M8).

DynaMut2 analyses are available under the following job IDs: 179128965804 (monomer); 179129026845 (8IM2); 179129312846 (multimer – HPD-TTC36).

