## Supplementarty data for "Tuning inhibitor susceptibility of human 4-hydroxyphenylpyruvate dioxygenase to rescue tyrosine catabolism in the presence of β-triketones"

**Supplementary information**

**HPD cDNA codon-optimized for *E. coli*:**

ATGACCACCTATAGCGATAAAGGTGCAAAACCGGAACGTGGTCGTTTTCTGCATTTTCATAGCGTTACCTTTTGGGTGGGTAATGCAAAACAGGCAGCAAGCTTTTATTGTAGCAAAATGGGTTTTGAACCGCTGGCATATCGTGGTCTGGAAACCGGTAGCCGTGAAGTTGTTAGCCATGTTATTAAACAGGGCAAAATCGTTTTTGTTCTGAGCAGCGCACTGAATCCGTGGAATAAAGAAATGGGTGATCATCTGGTTAAACATGGTGATGGCGTTAAAGATATTGCCTTTGAAGTTGAGGATTGCGATTACATTGTTCAGAAAGCACGTGAACGTGGTGCCAAAATTATGCGTGAACCGTGGGTTGAACAGGATAAATTTGGCAAAGTTAAATTTGCGGTGCTGCAGACCTATGGTGATACCACCCATACCCTGGTTGAAAAAATGAACTATATTGGCCAGTTTCTGCCTGGTTATGAAGCACCGGCATTTATGGATCCGCTGCTGCCGAAACTGCCTAAATGTAGCCTGGAAATGATTGATCATATTGTGGGTAATCAGCCGGATCAAGAAATGGTTAGCGCAAGCGAATGGTATCTGAAAAATCTGCAGTTTCACCGTTTTTGGAGCGTTGATGATACCCAGGTTCATACCGAATATAGCAGCCTGCGTAGCATTGTTGTTGCCAATTATGAAGAAAGCATCAAAATGCCGATTAATGAACCGGCACCGGGTAAAAAAAAAAGCCAGATTCAAGAGTATGTGGACTATAATGGTGGTGCCGGTGTTCAGCATATTGCACTGAAAACCGAAGATATTATCACCGCAATTCGTCATCTGCGTGAACGCGGTCTGGAATTTCTGAGCGTTCCGAGCACCTATTACAAACAGCTGCGTGAAAAACTGAAAACGGCCAAAATCAAAGTGAAAGAAAACATCGATGCACTGGAAGAACTGAAAATCCTGGTGGATTATGATGAGAAAGGTTATCTGCTGCAGATCTTTACCAAACCGGTTCAGGATCGTCCGACCCTGTTTCTGGAAGTTATTCAGCGTCATAATCATCAGGGTTTTGGTGCGGGTAACTTTAACAGCCTGTTTAAAGCATTTGAAGAGGAACAGAATCTGCGTGGTAA

**Table S1**: custom-designed phosphothioate (PTO)-primers for the HPD insert and vector backbone where nucleotide connecting phosphates contain a sulfur atom. The PTO nucleotides are located at the 5’-end of the primers.

| **Primer** | **Template** | **Length (bp)** | **Tm (°C)** |
| --- | --- | --- | --- |
| Fw insert (HPD) | 5’ – ctt taa gaa gga gat ata caT ATG ACC ACC TAT AGC - 3’ | 36 | 57.4 |
| Rv insert (HPD) | 5’ - tta gca gcc tag gtA TTA ATC AAT TAG TGG T - 3’ | 31 | 57.4 |
| Fw vector | 5’ - acc tag gct gct aaA CAA AGC - 3’ | 21 | 55.3 |
| Rv vector | 5’ - tgt ata tct cct tct taa agT TAA ACA AAA TTA TTT CTA G - 3’ | 40 | 55.0 |

**Table S2**: 22c-trick primers used for SSM at positions identified from Phase I of KnowVolution. For each targeted position, a primer pair was designed using the 22c-trick degenerate codon strategy to introduce the remaining 19 non-WT amino acids at the selected site in Phase II. Primer length, melting temperature (Tm; °C), and GC content (%) were calculated using the OligoAnalyzer tool. Degenerate bases are according to the IUPAC code: N = A/C/G/T; D = A/G/T; V = A/C/G; H = A/C/T; B = C/G/T. For each primer, the primer ID, melting temperature (Tm; °C) used in the PCR reaction, and sequence (5'→3') are provided.

| **Position** | **Primer** | **Template** | **Length (bp)** | **Tm (°C)** |
| --- | --- | --- | --- | --- |
| **M118** | Fw | 5'- GGT GCC AAA ATT NDT CGT GAA CCG TGG GT -3' | 29 | 62.9 – 66.2 |
|  |  | 5'- GGT GCC AAA ATT VHG CGT GAA CCG TGG GT -3' |  | 64.7 – 68.4 |
|  |  | 5'- GGT GCC AAA ATT TGG CGT GAA CCG TGG GT -3' |  | 66.8 |
|  | Rv | 5'- ACC CAC GGT TCA CGA HNA ATT TTG GCA CC -3' |  | 62.9 – 66.2 |
|  |  | 5'- ACC CAC GGT TCA CGC DBA ATT TTG GCA CC -3' |  | 64.7 – 68.4 |
|  |  | 5'- ACC CAC GGT TCA CGC CAA ATT TTG GCA CC -3' |  | 66.8 |
| **Q137** | Fw | 5'- TTG CGG TGC TGC NDT CCT ATG GTG ATA CC -3' | 29 | 63.2 – 66.8 |
|  |  | 5'- TTG CGG TGC TGC VHG CCT ATG GTG ATA CC -3' |  | 65.7 – 68.9 |
|  |  | 5'- TTG CGG TGC TGC TGG CCT ATG GTG ATA CC -3' |  | 67.1 |
|  | Rv | 5'- GGT ATC ACC ATA GGA HNG CAG CAC CGC AA -3' |  | 63.2 – 66.8 |
|  |  | 5'- GGT ATC ACC ATA GGC DBG CAG CAC CGC AA -3' |  | 65.7 – 68.9 |
|  |  | 5'- GGT ATC ACC ATA GGC CAG CAG CAC CGC AA -3' |  | 67.1 |
| **N151** | Fw | 5'- CTG GTT GAA AAA ATG NDT TAT ATT GGC CAG -3' | 30 | 55.6 – 58.9 |
|  |  | 5'- CTG GTT GAA AAA ATG VHG TAT ATT GGC CAG -3' |  | 56.8 – 60.2 |
|  |  | 5'- CTG GTT GAA AAA ATG TGG TAT ATT GGC CAG -3' |  | 58.4 |
|  | Rv | 5'- CTG GCC AAT ATA AHN CAT TTT TTC AAC CAG -3' |  | 55.6 – 58.9 |
|  |  | 5'- CTG GCC AAT ATA CDB CAT TTT TTC AAC CAG -3' |  | 56.8 – 60.2 |
|  |  | 5'- CTG GCC AAT ATA CCA CAT TTT TTC AAC CAG -3' |  | 58.4 |
| **I153** | Fw | 5'- GGT TGA AAA AAT GAA CTA TND TGG CCA GTT TCT GCC T -3' | 37 | 61.8 - 64.3 |
|  |  | 5'- GGT TGA AAA AAT GAA CTA TVH GGG CCA GTT TCT GCC T -3' |  | 62.8 - 65.6 |
|  |  | 5'- GGT TGA AAA AAT GAA CTA TTG GGG CCA GTT TCT GCC T -3' |  | 64.4 |
|  | Rv | 5'- AGG CAG AAA CTG GCC AHN ATA GTT CAT TTT TTC AAC C -3' |  | 61.8 - 64.3 |
|  |  | 5'- AGG CAG AAA CTG GCC CDB ATA GTT CAT TTT TTC AAC C -3' |  | 62.8 - 65.6 |
|  |  | 5'- AGG CAG AAA CTG GCC CCA ATA GTT CAT TTT TTC AAC C -3' |  | 64.4 |
| **M166** | Fw | 5'- GCA CCG GCA TTT NDT GAT CCG CTG CTG -3' | 27 | 64 – 67.6 |
|  |  | 5'- GCA CCG GCA TTT VHG GAT CCG CTG CTG -3' |  | 65.5 – 69.5 |
|  |  | 5'- GCA CCG GCA TTT TGG GAT CCG CTG CTG -3' |  | 67.7 |
|  | Rv | 5'- CAG CAG CGG ATC AHN AAA TGC CGG TGC -3' |  | 64 – 67.6 |
|  |  | 5'- CAG CAG CGG ATC CDB AAA TGC CGG TGC -3' |  | 65.5 – 69.5 |
|  |  | 5'- CAG CAG CGG ATC CCA AAA TGC CGG TGC -3' |  | 67.7 |
| **S177** | Fw | 5'- GAA ACT GCC TAA ATG TND TCT GGA AAT GAT TGA T -3' | 34 | 57.7 – 60.4 |
|  |  | 5'- GAA ACT GCC TAA ATG TVH GCT GGA AAT GAT TGA T -3' |  | 59.2 – 62.3 |
|  |  | 5'- GAA ACT GCC TAA ATG TTG GCT GGA AAT GAT TGA T -3' |  | 60.9 |
|  | Rv | 5'- ATC AAT CAT TTC CAG AHN ACA TTT AGG CAG TTT C -3' |  | 57.7 – 60.4 |
|  |  | 5'- ATC AAT CAT TTC CAG CDB ACA TTT AGG CAG TTT C -3' |  | 59.2 – 62.3 |
|  |  | 5'- ATC AAT CAT TTC CAG CCA ACA TTT AGG CAG TTT C -3' |  | 60.9 |
| **I278** | Fw | 5'- GAT ATT ATC ACC GCA NDT CGT CAT CTG CGT G -3' | 31 | 60.3 – 63.3 |
|  |  | 5'- GAT ATT ATC ACC GCA VHG CGT CAT CTG CGT G -3' |  | 62.4 – 65.2 |
|  |  | 5'- GAT ATT ATC ACC GCA TGG CGT CAT CTG CGT G -3' |  | 63.7 |
|  | Rv | 5'- CAC GCA GAT GAC GAH NTG CGG TGA TAA TAT C -3' |  | 60.3 – 63.3 |
|  |  | 5'- CAC GCA GAT GAC GCD BTG CGG TGA TAA TAT C -3' |  | 62.4 – 65.2 |
|  |  | 5'- CAC GCA GAT GAC GCC ATG CGG TGA TAA TAT C -3' |  | 63.7 |
| **K302** | Fw | 5'- CAG CTG CGT GAA NDT CTG AAA ACG GCC AA -3' | 29 | 62.8 – 66.1 |
|  |  | 5'- CAG CTG CGT GAA VHG CTG AAA ACG GCC AA -3' |  | 65.2 – 68.2 |
|  |  | 5'- CAG CTG CGT GAA TGG CTG AAA ACG GCC AA -3' |  | 66.6 |
|  | Rv | 5'- TTG GCC GTT TTC AGA HNT TCA CGC AGC TG -3' |  | 62.8 – 66.1 |
|  |  | 5'- TTG GCC GTT TTC AGC DBT TCA CGC AGC TG -3' |  | 65.2 – 68.2 |
|  |  | 5'- TTG GCC GTT TTC AGC CAT TCA CGC AGC TG -3' |  | 66.6 |
| **K307** | Fw | 5'- ACT GAA AAC GGC CND TAT CAA AGT GAA AG -3' | 29 | 57.3 – 60.9 |
|  |  | 5'- ACT GAA AAC GGC CVH GAT CAA AGT GAA AG -3' |  | 59.6 - 63 |
|  |  | 5'- ACT GAA AAC GGC CTG GAT CAA AGT GAA AG -3' |  | 61 |
|  | Rv | 5'- CTT TCA CTT TGA TAH NGG CCG TTT TCA GT -3' |  | 57.3 – 60.9 |
|  |  | 5'- CTT TCA CTT TGA TCD BGG CCG TTT TCA GT -3' |  | 59.6 - 63 |
|  |  | 5'- CTT TCA CTT TGA TCC AGG CCG TTT TCA GT -3' |  | 61 |

**Table S3**: 22c-trick primers used for SSM at positions identified from Phase III (molecular modelling) of KnowVolution. For each targeted position, a primer pair was designed using the 22c-trick degenerate codon strategy to introduce the remaining 19 non-WT amino acids at the selected site in Phase II. Primer length, melting temperature (Tm; °C), and GC content (%) were calculated using the OligoAnalyzer tool. Degenerate bases are according to the IUPAC code: N = A/C/G/T; D = A/G/T; V = A/C/G; H = A/C/T; B = C/G/T. For each primer, the primer ID, melting temperature (Tm; °C) used in the PCR reaction, and sequence (5'→3') are provided.

| **Position** | **Primer** | **Template** | **Length (bp)** | **Tm (°C)** |
| --- | --- | --- | --- | --- |
| **P239** | Fw | 5'- G AAA GCA TCA AAA TGN DTA TTA ATG AAC CGG -3' | 30 | 55.2 – 58.4 |
|  |  | 5'- G AAA GCA TCA AAA TGV HGA TTA ATG AAC CGG -3' |  | 56.7 – 60.1 |
|  |  | 5'- G AAA GCA TCA AAA TGT GGA TTA ATG AAC CGG -3' |  | 58.4 |
|  | Rv | 5'- CCG GTT CAT TAA TAH NCA TTT TGA TGC TTT C -3' |  | 55.2 – 58.4 |
|  |  | 5'- CCG GTT CAT TAA TCD BCA TTT TGA TGC TTT C -3' |  | 56.7 – 60.1 |
|  |  | 5'- CCG GTT CAT TAA TCC ACA TTT TGA TGC TTT C -3' |  | 58.4 |
| **N241** | Fw | 5'- GC ATC AAA ATG CCG ATT NDT GAA CCG GCA CC -3' | 29 | 61.5 – 64.8 |
|  |  | 5'- GC ATC AAA ATG CCG ATT VHG GAA CCG GCA CC -3' |  | 62.9 – 66.6 |
|  |  | 5'- GC ATC AAA ATG CCG ATT TGG GAA CCG GCA CC -3' |  | 64.9 |
|  | Rv | 5'- GGT GCC GGT TCA HNA ATC GGC ATT TTG ATG C -3' |  | 61.5 – 64.8 |
|  |  | 5'- GG TGC CGG TTC CDB AAT CGG CAT TTT GATGC -3' |  | 62.9 – 66.6 |
|  |  | 5'- GG TGC CGG TTC CCA AAT CGG CAT TTT GAT GC -3' |  | 64.9 |
| **Q334** | Fw | 5'- GG TTA TCT GCT GND TAT CTT TAC CAA ACC G -3' | 29 | 57.5 – 60.7 |
|  |  | 5'- GGT TAT CTG CTG VHG ATC TTT ACC AAA CCG -3' |  | 59 – 62.4 |
|  |  | 5'- GGT TAT CTG CTG TGG ATC TTT ACC AAA CCG -3' |  | 60.7 |
|  | Rv | 5'- CGG TTT GGT AAA GAT AHN CAG CAG ATA ACC -3' |  | 57.5 – 60.7 |
|  |  | 5'- CGG TTT GGT AAA GAT CDB CAG CAG ATA ACC -3' |  | 59 – 62.4 |
|  |  | 5'- CGG TTT GGT AAA GAT CCA CAG CAG ATA ACC -3' |  | 60.7 |
| **N363** | Fw | 5'- GGT TTT GGT GCG GGT NDT TTT AAC AGC CTG T-3' | 31 | 60.6 – 63.7 |
|  |  | 5'- GGT TTT GGT GCG GGT VHG TTT AAC AGC CTG T -3' |  | 61.8 – 65.3 |
|  |  | 5'- GGT TTT GGT GCG GGT TGG TTT AAC AGC CTG T -3' |  | 63.7 |
|  | Rv | 5'- ACA GGC TGT TAA AAH NAC CCG CAC CAA AAC C -3' |  | 60.6 – 63.7 |
|  |  | 5'- ACA GGC TGT TAA ACD BAC CCG CAC CAA AAC C -3' |  | 61.8 – 65.3 |
|  |  | 5'- ACA GGC TGT TAA ACC AAC CCG CAC CAA AAC C -3' |  | 63.7 |
| **F364** | Fw | 5'- GGT GCG GGT AAC NDT AAC AGC CTG TTT AAA GC-3' | 32 | 62.1 – 65.2 |
|  |  | 5'- GGT GCG GGT AAC VHG AAC AGC CTG TTT AAA GC-3' |  | 64.1 – 67 |
|  |  | 5'- GGT GCG GGT AAC TGG AAC AGC CTG TTT AAA GC -3' |  | 65.3 |
|  | Rv | 5'- GCT TTA AAC AGG CTG TTA HNG TTA CCC GCA CC -3' |  | 62.1 – 65.2 |
|  |  | 5'- GCT TTA AAC AGG CTG TTC DBG TTA CCC GCA CC -3' |  | 64.1 – 67 |
|  |  | 5'- GCT TTA AAC AGG CTG TTC CAG TTA CCC GCA CC -3' |  | 65.3 |
| **L367** | Fw | 5'- GCG GGT AAC TTT AAC AGC NDT TTT AAA GCA TTT G -3' | 34 | 58.7 – 61.6 |
|  |  | 5'- GCG GGT AAC TTT AAC AGC VHG TTT AAA GCA TTT G -3' |  | 60.3 – 63 |
|  |  | 5'- GCG GGT AAC TTT AAC AGC TGG TTT AAA GCA TTT G -3' |  | 61.4 |
|  | Rv | 5'- CAA ATG CTT TAA AAH NGC TGT TAA AGT TAC CCG C -3' |  | 58.7 – 61.6 |
|  |  | 5'- CAA ATG CTT TAA ACD BGC TGT TAA AGT TAC CCG C -3' |  | 60.3 – 63 |
|  |  | 5'- CAA ATG CTT TAA ACC AGC TGT TAA AGT TAC CCG C -3' |  | 61.4 |

**Table S4**: Primers designed for recombination introduced by SDM. Primer length, melting temperature (Tm; °C), and GC content (%) were calculated using the OligoAnalyzer tool. The red colored codon indicates the site of the introduced substitution.

| **Position** | **Substitution** | **Primer** | **Template** | **Length (bp)** | **Tm (°C)** |
| --- | --- | --- | --- | --- | --- |
| 118 | M 🡪 I | Fw  Rv | 5'- GGT GCC AAA ATT ATT CGT GAA CCG TGG -3'  5'- CCA CGG TTC ACG AAT AAT TTT GGC ACC -3' | 27 | 60.5 |
| 138 | T 🡪 S | Fw | 5'- TTG CGG TGC TGC AGT CCT ATG GTG ATA CC -3'  5'- GGT ATC ACC ATA GGA CTG CAG CAC CGC AA -3' | 29 | 65.1 |
|  |  | Rv |  |  |  |
| 151 & 153 | N 🡪 D | Fw | 5’ - GTT GAA AAA ATG GAT TAT GAG GGC CAG TTT CTG C – 3’  5’ - G CAG AAA CTG GCC CTC ATA ATC CAT TTT TTC AAC -3' | 34 | 61.3 |
|  | I 🡪 E | Rv |  |  |  |
| 177 | S 🡪 Q | Fw | 5'- GAA ACT GCC TAA ATG TCA GCT GGA AAT GAT TG -3'  5'- C AAT CAT TTC CAG CTG ACA TTT AGG CAG TTT C -3' | 32 | 59.8 |
|  |  | Rv |  |  |  |
| 278 | I 🡪 V | Fw | 5'- GAT ATT ATC ACC GCA GTT CGT CAT CTG CGT G -3'  5'- CAC GCA GAT GAC GAA CTG CGG TGA TAA TAT C -3' | 31 | 61.9 |
|  |  | Rv |  |  |  |
| 302 & 307 | K 🡪 R | Fw  Rv | 5'- CTG CGT GAA CGT CTG AAA ACG GCC AAT ATC AAA GTG -3'  5'- CAC TTT GAT ATT GGC CGT TTT CAG ACG TTC ACG CAG -3' | 36 | 64.6 |

**Table S5**: DynaMut2 predicted stability change (ΔΔG^Stability^) of single HPD mutants *versus* HPD-WT (kcal/mol).

| **mutation** | **monomer** | **I8M2** | **multimer** | **average** | **sd** |
| --- | --- | --- | --- | --- | --- |
| M118I | -0.52 | -0.47 | -0.51 | -0.50 | 0.03 |
| M118L | -0.68 | -0.66 | -0.69 | -0.68 | 0.02 |
| Q137R | -0.39 | -0.35 | -0.67 | -0.47 | 0.17 |
| T138S | -0.48 | -0.66 | -0.4 | -0.51 | 0.13 |
| N151D | 0.31 | 0.57 | 0.27 | 0.38 | 0.16 |
| I153D | 0.15 | 0.37 | 0.19 | 0.24 | 0.12 |
| I153E | 0.66 | 0.73 | 0.7 | 0.70 | 0.04 |
| I153N | 0 | 0.07 | 0.13 | 0.07 | 0.07 |
| M166S | 0.67 | 0.7 | 0.79 | 0.72 | 0.06 |
| M166T | 0.54 | 0.72 | 0.74 | 0.67 | 0.11 |
| S177G | -0.74 | -0.3 | -0.3 | -0.45 | 0.25 |
| S177Q | -0.82 | -0.74 | -0.72 | -0.76 | 0.05 |
| I278V | -1.2 | -2 | -1.15 | -1.45 | 0.48 |
| K302R | -0.33 | -0.52 | -0.41 | -0.42 | 0.10 |
| K307N | 0.06 | 0.21 | 0.07 | 0.11 | 0.08 |
| K307P | 0.41 | 0.4 | 0.43 | 0.41 | 0.02 |

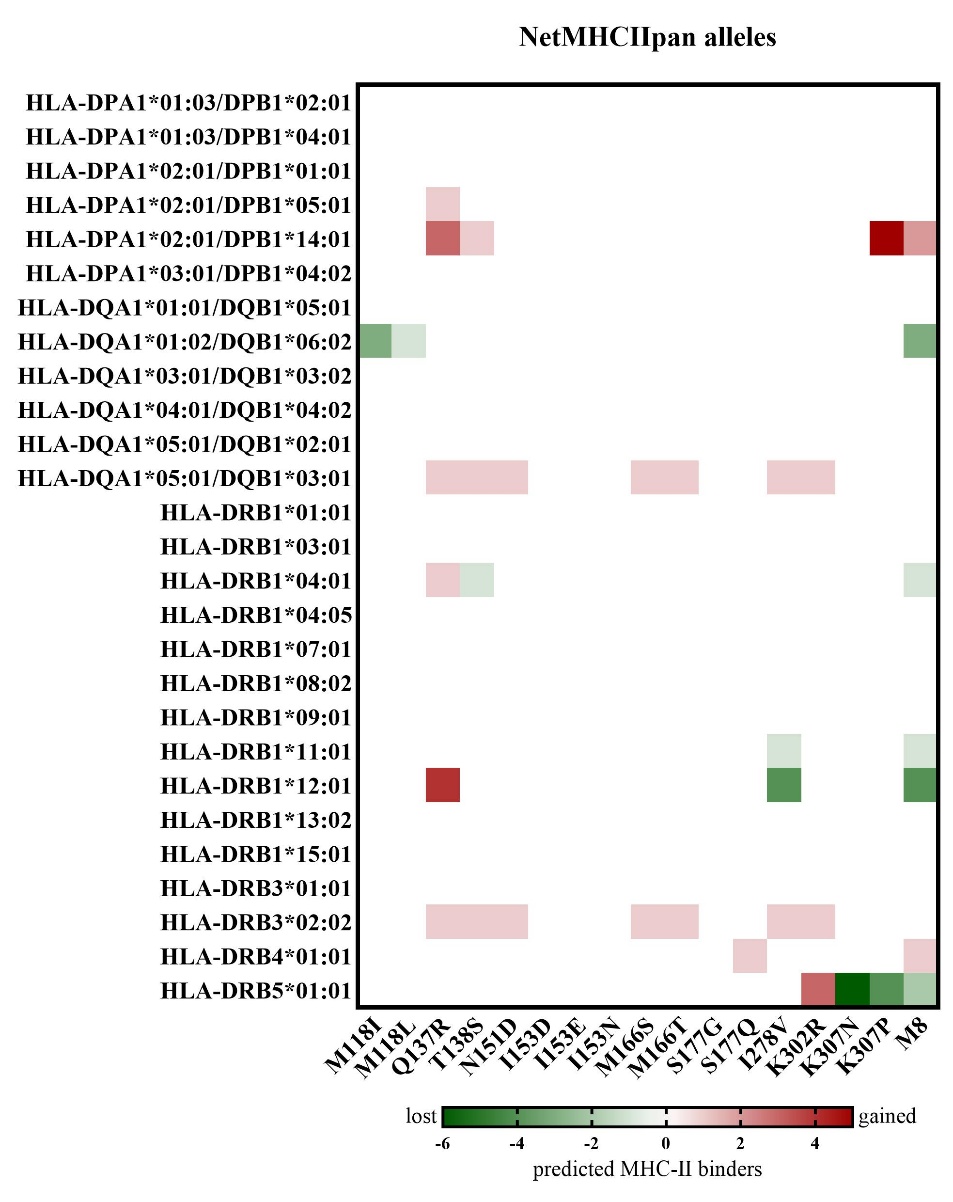
**Fig. S1**: *In silico* analysis of the effects of identified HPD substitutions on predicted MHC-II binding alleles. Changes in the number of predicted MHC-II binders for specific alleles determined using NetMHCIIpan. Values represent the net change in predicted binders relative to HPD-WT, with green indicating a loss and red indicating a gain of predicted MHC-II binders. M8 represents the lead variant containing eight accumulated substitutions.
